# DAF-18/PTEN protects LIN-35/Rb from CLP-1/CAPN-mediated cleavage to promote starvation resistance

**DOI:** 10.1101/2025.02.17.638677

**Authors:** Jingxian Chen, Rojin Chitrakar, L. Ryan Baugh

## Abstract

Starvation resistance is a fundamental trait with profound influence on fitness and disease risk. DAF-18, the *C. elegans* ortholog of the tumor suppressor PTEN, promotes starvation resistance. PTEN is a dual phosphatase, and DAF-18 promotes starvation resistance as a lipid phosphatase by antagonizing insulin/IGF and PI3K signaling, activating the tumor suppressor DAF-16/FoxO. However, if or how DAF-18/PTEN protein-phosphatase activity promotes starvation resistance is unknown. Using genetic, genomic, bioinformatic, and biochemical approaches, we identified the *C. elegans* retinoblastoma/RB protein homolog, LIN-35/Rb, as a critical mediator of the effect of DAF-18/PTEN on starvation resistance. We show that DAF-18/PTEN protects LIN-35/Rb from cleavage by the μ-Calpain homolog CLP-1/CAPN, and that LIN-35/Rb together with the repressive DREAM complex promote starvation resistance. We conclude that the tumor suppressors DAF-18/PTEN and LIN-35/Rb function in a linear pathway, with LIN-35/Rb and the rest of the DREAM complex functioning as a transcriptional effector of DAF-18/PTEN protein-phosphatase activity resulting in repression of germline gene expression. This work is significant for revealing a network of tumor suppressors that promote survival during cellular and developmental quiescence.

## INTRODUCTION

One of the most fascinating things about biology is the ability of organisms to robustly adapt to different environmental conditions. Many animals enter developmental diapause or a diapause-like state to endure unfavorable conditions (Easwaran & Montell, 2023). In the nematode *Caenorhabditis elegans*, when embryos hatch in the complete absence of food, they arrest development in the first larval stage (L1 arrest or L1 diapause) (Baugh, 2013). There is no cell proliferation, migration or fusion during L1 arrest (Baugh & Sternberg, 2006), and gene expression and metabolism are dramatically altered to support survival (Baugh *et al*, 2009; Hibshman *et al*, 2017; Webster *et al*, 2022). Larvae can survive L1 arrest for weeks (Johnson *et al*, 1984), they continue foraging, and they recover upon feeding, albeit with developmental delay (Olmedo *et al*, 2020) and reproductive costs (Jobson *et al*, 2015; Jordan *et al*, 2019) commensurate with the amount of time spent in arrest.

How long worms survive L1 arrest is regulated by a variety of conserved signaling pathways and processes, with insulin/insulin-like growth factor signaling (IIS) being critical (Baugh & Hu, 2020). During starvation, IIS is reduced, with activity of DAF-2/insulin-like growth factor receptor (IGFR) and downstream AGE-1/phosphoinositide 3-kinase (PI3K) signaling decreased. DAF-2/IGFR and AGE-1/PI3K antagonize the Forkhead box O (FoxO) transcription factor DAF-16 (Lin *et al*, 1997; Ogg *et al*, 1997), and DAF-16/FoxO nuclear localization and activity increase during L1 arrest (Hibshman *et al*., 2017; Weinkove *et al*, 2006). Nuclear DAF-16/FoxO activates transcription of genes that promote starvation resistance (Hibshman *et al*., 2017) and represses genes that promote postembryonic development (Kaplan *et al*, 2015). Consequently, loss of *daf-16* renders worms starvation-sensitive, with compromised survival (Munoz & Riddle, 2003), and arrest-defective, with postembryonic development initiated in starved L1 larvae (Baugh & Sternberg, 2006). Mammalian FoxO proteins are tumor suppressors (Paik *et al*, 2007), and *daf-16/FoxO* suppresses tumors in *C. elegans* (Pinkston *et al*, 2006). Despite substantial impacts on phenotype, *daf-16/FoxO* does not account for much of the transcriptional response to starvation (Hibshman *et al*., 2017), suggesting IIS-independent regulation of transcription.

The tumor suppressor phosphatase and tensin (PTEN) is a potent negative regulator of IIS in humans and *C. elegans* (Chalhoub & Baker, 2009; Murphy & Hu, 2013). PTEN was originally called MMAC, because it is mutated in multiple advanced cancers at high frequency (Li *et al*, 1997; Steck *et al*, 1997). The sole *C. elegans* PTEN homolog DAF-18 is required for L1 arrest, and *daf-18/PTEN* mutants are arrest-defective (Fukuyama *et al*, 2006) and starvation-sensitive (Fukuyama *et al*, 2012), like *daf-16* mutants, though more severe in both cases. *daf-18/PTEN* is also required to repress germline gene expression during L1 arrest (Fry *et al*, 2021), but it is unclear how it exerts transcriptional repression. PTEN is best known for its lipid-phosphatase activity, which inhibits PI3K signaling by converting phosphatidylinositol-3, 4, 5-triphosphate (PIP3) to phosphatidylinositol-4, 5-bisphosphate (PIP2) in mammals and *C. elegans* (Myers *et al*, 1998; Ogg & Ruvkun, 1998). However, PTEN is also a protein-phosphatase (Myers & Tonks, 1997), and PTEN protein-phosphatase substrates with different roles in signaling and cell migration have been identified in mammals (Abbas *et al*, 2019; Gu *et al*, 1999; Gu *et al*, 2011; Shi *et al*, 2014; Tamura *et al*, 1998). Point mutations intended to selectively disrupt the lipid and protein-phosphatase activities of DAF-18/PTEN have been reported (Brisbin *et al*, 2009; Nakdimon *et al*, 2012; Solari *et al*, 2005; Zheng *et al*, 2018), supposedly providing a genetic approach to dissect protein function. However, these two alleles fail to complement each other for dauer formation and L1 starvation-resistance phenotypes, indicating that they do not selectively disrupt these two activities (Chen *et al*, 2022; Wittes & Greenwald, 2022). It remains unclear whether DAF-18/PTEN functions as a protein phosphatase in *C. elegans*, if such activity contributes to regulation of L1 arrest, and, if so, how.

Like DAF-18/PTEN, LIN-35/Rb is a tumor-suppressor homolog that promotes starvation resistance in starved L1 larvae (Cui *et al*, 2013). *lin-35/Rb* mutants also have a synthetic multivulva (Muv) phenotype (Lu & Horvitz, 1998) and express germline genes in the soma (Petrella *et al*, 2011; Rechtsteiner *et al*, 2019; Wang *et al*, 2005; Wu *et al*, 2012). LIN-35 is the sole *C. elegans* homolog of the human retinoblastoma (RB) pocket protein family (Lu & Horvitz, 1998). There are three RB-encoding genes in mammals, *RB1*, *RBL1*, and *RBL2* (Henley & Dick, 2012). *RB1* was the first tumor suppressor to be cloned and characterized (Berry *et al*, 2019), and it was later found to be defective in many human cancers in addition to retinoblastoma (Burkhart & Sage, 2008). All three RB pocket proteins bind to adenovirus early region 2 binding factor (E2F) family transcription factors and the E2F dimerizing partner (DP) family members (Henley & Dick, 2012). Together with E2F, the protein encoded by *RB1*, pRB, enforces the G1/S cell cycle checkpoint through transcriptional repression (Uchida, 2016). In addition to E2F, the protein products of *RBL1* and *RBL2* (p107 and p130, respectively) recruit MuvB proteins (Guiley *et al*, 2015), and DP, RB, and E2F family proteins plus five MuvB proteins form the DREAM complex, which mediates transcriptional repression (Fischer *et al*, 2022). Mutation of *C. elegans lin-37*, which encodes a MuvB/DREAM protein, results in ectopic expression of germline genes in the soma, like *lin-35/Rb* (Petrella *et al*., 2011). However, because LIN-35 is the sole pocket protein in *C. elegans*, it is unclear whether it regulates starvation resistance independent of the MuvB complex like pRB, as a component of the DREAM complex like p107 and p130, or via another mechanism.

DAF-18/PTEN and LIN-35/Rb are both important regulators of starvation resistance in *C. elegans*. However, whether there is a functional link between them has not been addressed, nor has a connection between these two paramount tumor suppressors been reported in mammals. Here, we show that DAF-18/PTEN promotes starvation resistance independently of IIS, likely through its protein-phosphatase activity. Using a combination of genetics, functional genomics, and biochemistry, we discovered that *daf-18/PTEN* regulates LIN-35/Rb at the protein level to promote starvation resistance. LIN-35/Rb is cleaved and destabilized in the absence of *daf-18/PTEN*. We show that the human μ-Calpain protease homolog CLP-1/CAPN is responsible for this negative regulation of LIN-35/Rb and that *daf-18/PTEN* is a negative regulator of CLP-1/CAPN. We also report that the DREAM complex functions downstream of DAF-18/PTEN via LIN-35/Rb to promote starvation resistance likely by repressing germline gene expression. Our results provide a significant functional connection between DAF-18/PTEN and LIN-35/Rb that is likely conserved, and they identify a transcriptional effector mechanism of DAF-18/PTEN protein-phosphatase activity. Our insights into how these tumor suppressors promote survival during developmental quiescence have important implications for cancer, stem cell maintenance, and organismal fitness during starvation.

## RESULTS

### *daf-18/PTEN* promotes starvation resistance independently of AGE-1/PI3K signaling and ***daf-16/FoxO***

We used genetic analysis to examine whether DAF-18/PTEN functions independently of AGE-1/PI3K signaling to regulate L1 starvation resistance. In contrast to *daf-18/PTEN* (Baugh & Sternberg, 2006; Fukuyama *et al*., 2012), mutation of *age-1/PI3K* increases L1 starvation resistance (Baugh & Sternberg, 2006; Cui *et al*., 2013; Fukuyama *et al*., 2012; Munoz & Riddle, 2003). There is no detectable PIP3 in *age-1* null mutants (Bharill *et al*, 2013), so we reasoned that if the only function of DAF-18/PTEN in this context is to counteract AGE-1/PI3K signaling by dephosphorylating PIP3 to produce PIP2 (as opposed to relying on protein-phosphatase activity), then loss of *age-1* should rescue resistance of an otherwise starvation-sensitive *daf-18* null mutant to wild-type levels or greater. However, the double null mutant *age-1(m333); daf-18(ok480)* (Brisbin *et al*., 2009; Morris *et al*, 1996) was significantly more starvation-sensitive than wild type (Fig 1A), suggesting that DAF-18/PTEN promotes starvation resistance independently of AGE-1/PI3K signaling.

**Fig 1.**
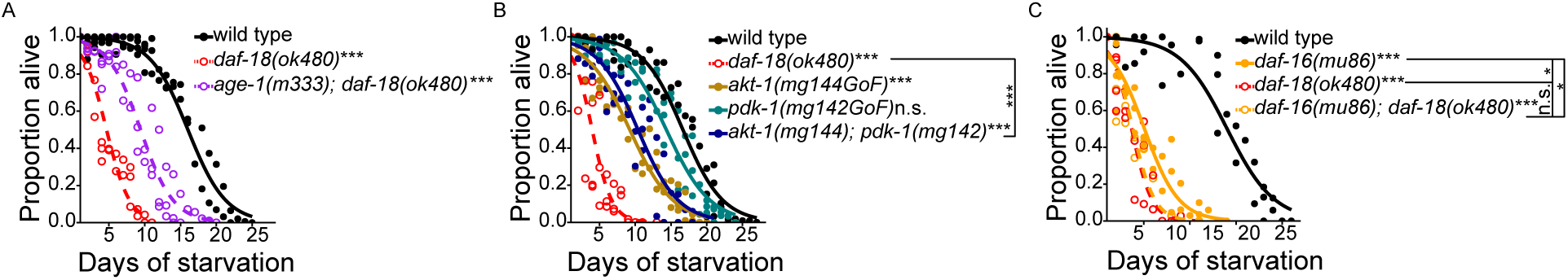
*daf-18/PTEN* promotes starvation resistance independently of AGE-1/PI3K signaling and *daf-16/FoxO*. (A-C) Proportion of survivors is plotted throughout L1 starvation. Survival was scored daily. Four biological replicates were performed. Two-tailed, unpaired, variance-pooled t-tests were performed on half-lives to compare genotypes. Unless otherwise noted with brackets, all comparisons were against wild type. *P < 0.05; ***P < 0.001; n.s. not significant. For details on the statistical method, see *Statistics for starvation survival* in Materials and Methods. (A) Number of scored animals was 105 +/- 25 (mean +/- standard deviation). (B) Number of scored animals was 104 +/- 31 (mean +/- standard deviation). (C) Number of scored animals was 105 +/- 27 (mean +/- standard deviation). Half-lives of all four genotypes were subjected to Levene’s test to assess variance homogeneity across groups, which suggested homogeneous variance. Half-lives were used in a two-way ANOVA (formula: half-life ∼ loss of *daf-18* * loss of *daf-16*) to assess additivity of the effects of *daf-18* and *daf-16* on survival, which yielded an interaction p-value of 7 x 10^-6^, indicating non-additivity (interaction or dependence) of *daf-18* and *daf-16*.

We assayed *akt-1/AKT* and *pdk-1/PDK* gain-of-function mutations to further examine AGE-1/PI3K-independent effects of DAF-18/PTEN. AGE-1/PI3K signaling activates PDK-1 and AKT-1 kinases, which antagonize DAF-16/FoxO effector function (Murphy & Hu, 2013). Both *pdk-1(mg142)* and *akt-1(mg144)* are strong gain-of-function alleles that completely abolish the *age-1* dauer-constitutive phenotype at 20°C (Paradis *et al*, 1999). We reasoned that if the only function of DAF-18/PTEN in promoting starvation resistance is to oppose AGE-1/PI3K signaling, then sufficiently increasing AGE-1/PI3K signaling activity should phenocopy *daf-18* null mutants. *akt-1(mg144)* significantly reduced starvation resistance, as expected, but not to the extent of the *daf-18* null allele *ok480* (Fig 1B)*. pdk-1(mg142)* had no significant effect on its own, and the *pdk-1(mg142); akt-1(mg144)* double mutant was no more sensitive than *akt-1(mg144)* alone. Notably, the double mutant did not phenocopy *daf-18(ok480)*, further suggesting that DAF-18/PTEN promotes starvation resistance independently of AGE-1/PI3K signaling, potentially through its putative protein-phosphatase activity.

We used genetic epistasis analysis to examine independence of *daf-16/FoxO* and *daf-18/PTEN* function. *daf-16/FoxO* is required for the *daf-2/IGFR* starvation-resistance phenotype (Baugh & Sternberg, 2006), suggesting it is the primary effector of AGE-1/PI3K signaling. However, a *daf-18* null mutant appears to be more sensitive to starvation than a *daf-16* null mutant (Baugh & Sternberg, 2006; Fukuyama *et al*., 2012), suggesting that loss of DAF-18 activity does more than decrease DAF-16 activity via increased PI3K signaling. We reproduced this observation, and the null allele *daf-18(ok480)* is significantly more starvation sensitive than the null allele *daf-16(mu86)* (Fig 1C). Notably, the double mutant was not different from *daf-18(ok480)* alone, showing that loss of *daf-16* does nothing in a *daf-18* null background. Indeed, a two-way ANOVA revealed a significant interaction between *daf-18* and *daf-16*, indicating non-additivity of the two mutations. These results are consistent with DAF-18/PTEN inhibiting PI3K/AKT signaling to activate DAF-16/FoxO, but they also support the conclusion that DAF-18/PTEN has an additional, independent function that promotes starvation resistance.

### Transcriptome-based epistasis analysis suggests *lin-35/Rb* mediates *daf-16/FoxO*-independent effects of *daf-18/PTEN*

We used mRNA sequencing (RNA-seq) to extend epistasis analysis of *daf-16/FoxO* and *daf-18/PTEN* to the transcriptome to isolate *daf-16*-independent effects of *daf-18* on gene expression. We performed bulk RNA-seq on *daf-16(mu86)* and *daf-18(ok480)* single null mutants, the *daf-16(mu86); daf-18(ok480)* double mutant, and wild type in starved L1 larvae (S1 Data). We wanted to capture relatively early, direct effects of *daf-16* and *daf-18* rather than indirect effects. We analyzed hatching in our staged populations and determined the timepoint when hatching first reached its maximum for all genotypes, and we collected our RNA-seq samples at that timepoint (S1 Fig A; for details, see *Hatching efficiency for determining RNA-seq sample collection timepoint* in Materials and Methods). This analysis suggests that each population was only ∼4 hr into L1 starvation on average, which is relatively early compared to the peak of the starvation response at about 12 hr (Baugh *et al*., 2009; Webster *et al*., 2022). Principal Component Analysis (PCA) of the RNA-seq data shows that all three mutants separate from wild type (S1 Fig B), suggesting relatively robust effects on gene expression despite the samples being collected early in starvation.

We used cluster analysis to get an overview of the RNA-seq results. We identified 871 genes that were differentially expressed across the four genotypes out of 15,018 detected genes, and we subjected the differentially expressed genes to hierarchical clustering (Fig 2A). Consistent with PCA (S1 Fig B), the three mutants clustered together and wild type stood alone in the dendrogram (Fig 2A). Consistent with their starvation-resistance phenotypes (Fig 1C), the expression profiles of *daf-18(ok480)* and *daf-16(mu86); daf-18(ok480)* were more similar to each other than *daf-16(mu86)* (Fig 2B). *daf-16* and *daf-18* appear to affect many genes in common, but there also appear to be many genes affected by *daf-18* but not *daf-16*, consistent with *daf-18* functioning independently of PI3K signaling and *daf-16*.

**Fig 2.**
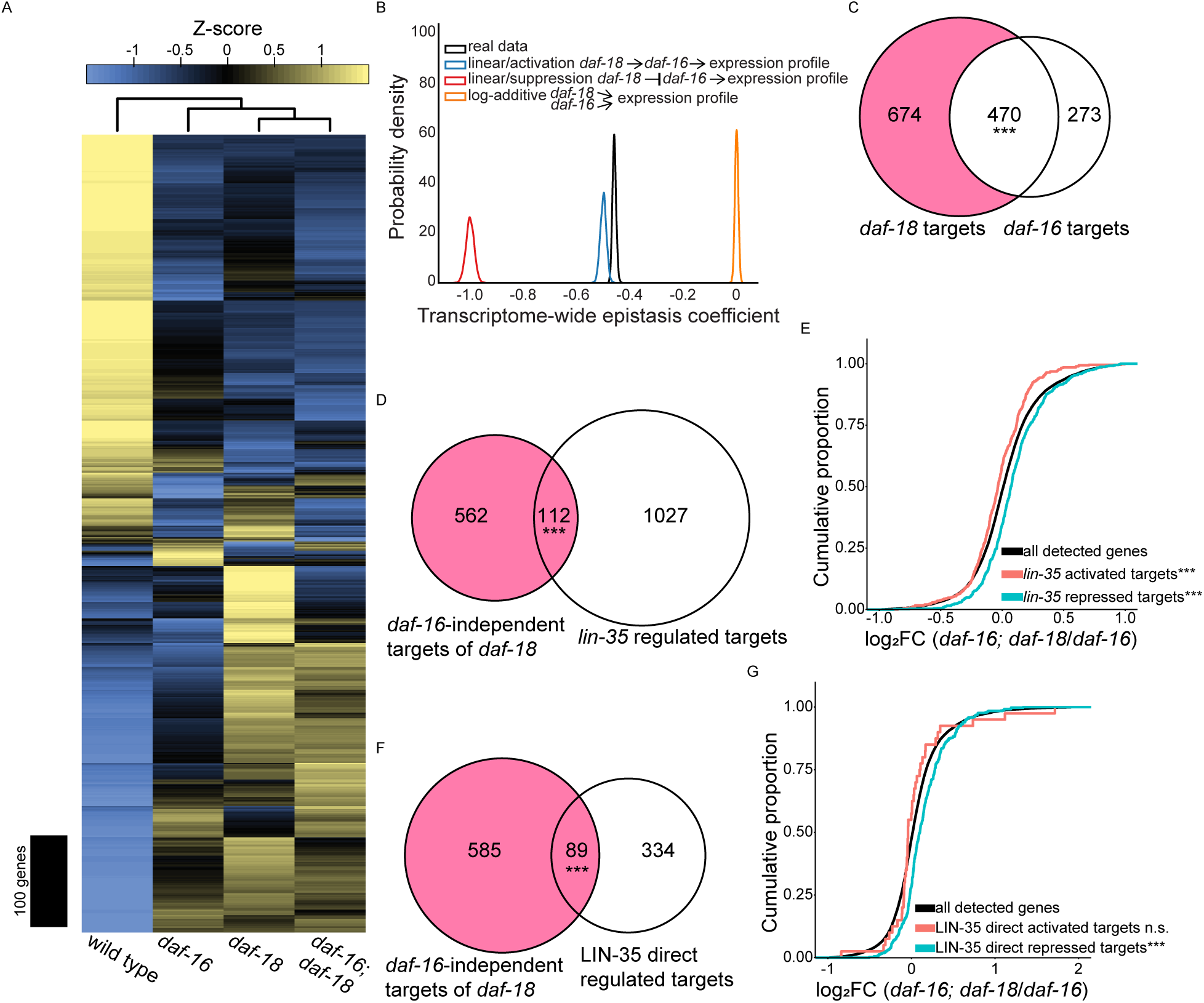
Transcriptome-based epistasis analysis suggests *lin-35/Rb* mediates *daf-16/FoxO*-independent effects of *daf-18/PTEN*. (A) RNA-seq heatmap of genes whose expression was significantly different across wild type, *daf-16(mu86)*, *daf-18(ok480)*, and *daf-16(mu86); daf-18(ok480)*. A generalized linear model found 871 genes out of 15,018 detected that were differentially expressed in the four genotypes tested. These 871 genes were z-score normalized and hierarchically clustered. Above the heatmap is the genotype dendrogram reflecting pairwise correlations between genotypes. For details on how these 871 genes were identified, see *Differential expression analysis of RNA-seq data* in Materials and Methods. (B) Distributions of transcriptome-wide epistasis coefficients are presented. RNA-seq data from wild type, *daf-16(mu86)* and *daf-18(ok480)* single mutants, and *daf-16(mu86); daf-18(ok480)* double mutant were bootstrapped to simulate transcriptome-wide epistasis coefficients (defined in (Angeles-Albores *et al*., 2018)) under three predefined null models for the regulatory relationship between *daf-18* and *daf-16*. Data were also bootstrapped to determine transcriptome-wide epistasis coefficient under a parameter-free model for the real data. Odds ratios (ORs) were calculated by dividing the likelihood of the parameter-free model by each predefined null model. Model rejection: OR > 10^3^. Linear/activation model OR: 2 x 10^4^; linear/suppression model OR: infinity; additive model OR: infinity. All three null models were rejected. For details on the statistics, see *Transcriptome-wide epistasis analysis* in Materials in Methods. (C) Overlap of differentially expressed genes (DEGs) in *daf-18* vs. wild type and *daf-16* vs. wild type during L1 arrest. (D) Overlap between *daf-16*-independent targets of *daf-18* and *lin-35/Rb* targets in L1 arrest (from expression analysis in (Cui *et al*., 2013)). (E) *lin-35* target gene (Cui *et al*., 2013) expression changes in *daf-16; daf-18* vs. *daf-16* are presented in cumulative distributions of log_2_ fold-change (FC). *lin-35* activated and repressed targets (n = 488 and n = 651, respectively) are genes whose expression decreased or increased in the *lin-35* mutant compared to wild type, respectively. (F) Overlap between *daf-16*-independent targets of *daf-18* and *lin-35* direct regulated targets in L1 arrest (determined by CHIP-seq and RNA-seq in (Gal *et al*., 2021); *i.e.*, genes bound by LIN-35 (direct) and whose expression was affected in the *lin-35* mutant (regulated)). (G) *lin-35* direct regulated targets (Gal *et al*., 2021) gene expression changes in *daf-16; daf-18* vs. *daf-16* are presented in cumulative distributions of log_2_ fold-change (FC). N = 40 for LIN-35 direct activated targets. N = 383 for LIN-35 direct repressed targets. (C, D and F) The pink region represents *daf-16*-independent targets of *daf-18*. Hypergeometric tests were performed to assess the overlap significance between two gene sets, with the background being all detected genes in RNA-seq (see S1 Data). (E and G) Kolmogorov-Smirnov tests were used to assess the equality between two cumulative distributions. All comparisons were against “all detected genes.” (C-G) ***P < 0.001; n.s. not significant. See also S1 Fig.

We performed a transcriptome-wide epistasis analysis (Angeles-Albores *et al*, 2018) to formalize our interpretations of the RNA-seq data. The rationale for this analysis is to use the results for each single mutant compared to wild type to generate expected results for the double mutant, assuming the mutants affect gene expression independently (log-additively). This is done for the 563 genes that are significantly differentially expressed in all three mutant genotypes compared to wild type, and bootstrapping and regression are used to determine a transcriptome-wide epistasis coefficient. We also simulated epistasis coefficients with our RNA-seq data using three predefined models for the relationship between *daf-18* and *daf-16*: linear/activation model where *daf-18* activates *daf-16*, and they are in a linear unbranched pathway; linear/suppression model where *daf-18* suppresses *daf-16*, and they are in a linear unbranched pathway; and a log-additive model where *daf-18* and *daf-16* act additively and independently of each other (in parallel). We compared the observed distribution of epistasis coefficients generated with the parameter-free model to the three distributions resulting from the predefined models, computed model likelihoods with Bayesian statistics, and calculated odds ratios for the simulation results for each predefined model compared to the observed coefficients. Both the linear/suppression model and the log-additive model had an odds ratio of positive infinity, suggesting they are highly unlikely to represent the real relationship between *daf-18* and *daf-16* (Fig 2B). This is consistent with DAF-18/PTEN inhibiting AGE-1/PI3K signaling to activate DAF-16/FoxO. However, the linear/activation model was rejected with an odds ratio of 2 x 10^4^ (Fig 2B), which suggests that *daf-18/PTEN* affects transcriptional regulation by doing more than activating DAF-16/FoxO.

Differential gene expression analysis also suggested *daf-16/FoxO*-independent effects of *daf-18/PTEN.* We identified significantly differentially expressed genes (DEGs) for each pair of genotypes (S1 Data). *daf-16/FoxO* and *daf-18/PTEN* shared a significant number DEGs compared to wild type (Fig 2C), as expected. However, *daf-18* had even more DEGs not in common with *daf-16* (“*daf-16*-independent targets of *daf-18*” highlighted in pink in Fig 2), suggesting a transcriptional effector in addition to DAF-16/FoxO.

We subjected the *daf-16*-independent targets of *daf-18* to gene set enrichment analysis (GSEA) using WormExp, which compares a user-supplied gene set to a database of gene sets defined by published functional genomic experiments (*e.g.*, RNA-seq with perturbation, ChIP-seq of specific proteins)(Yang *et al*, 2016). We found that genes from multiple experiments with *lin-35/Rb* perturbation were highly enriched (S2 Data). This was unexpected but not surprising since *lin-35/Rb* is known to promote L1 starvation resistance (Cui *et al*., 2013). We compared our *daf-16*-independent targets of *daf-18* to a set of “*lin-35* regulated targets” identified during L1 arrest (genes whose expression was affected by *lin-35* mutation in starved L1 larvae)(Cui *et al*., 2013), and confirmed significant overlap between these two gene sets (Fig 2D). These results reveal overlapping effects of *daf-18* (independent of PI3K signaling and *daf-16/FoxO*) and *lin-35/Rb* on gene expression during L1 arrest.

GSEA requires discrete gene sets, but we isolated *daf-16*-independent effects of *daf-18* for all genes by comparing *daf-16(mu86); daf-18(ok480)* and *daf-16(mu86)* and plotting cumulative distributions of log_2_ fold changes (log_2_FCs). “*Iin-35*-activated targets” (genes downregulated in *lin-35* mutant, starved L1 larvae compared to wild type) (Cui *et al*., 2013) had significantly smaller log_2_FCs compared to all detected genes (Fig 2E). Conversely, “*lin-35*-repressed targets” (genes upregulated in *lin-35* mutant, starved L1 larvae compared to wild type) (Cui *et al*., 2013) had significantly larger log_2_FCs. We also compared our *daf-16*-independent targets of *daf-18* to “LIN-35 direct regulated targets” (activated or repressed; determined by RNA-seq and ChIP-seq in starved L1 larvae) (Gal *et al*, 2021), and found significant overlap (Fig 2F). “LIN-35 direct repressed targets” had significantly larger log_2_FCs (*daf-16; daf-18*/*daf-16*) compared to all detected genes, while “LIN-35 direct activated targets” were indistinguishable (Fig 2G). As a negative control, we performed the same GSEA between both *lin-35* target sets (Cui *et al*., 2013; Gal *et al*., 2021) and our “*daf-18*-independent targets of *daf-16*” (the 273 genes in Fig 2C), and neither was significantly enriched (S1 Fig C and D). Our RNA-seq analysis suggests that *lin-35/Rb* and *daf-16/FoxO* independently affect gene expression during L1 arrest, and that *daf-18/PTEN* and *lin-35/Rb* function in a linear pathway, or possibly in parallel with convergence on a common set of regulatory targets.

### *lin-35/Rb* functions downstream of *daf-18/PTEN* to promote starvation resistance

We used epistasis analysis to determine if *daf-16/FoxO* and *lin-35/Rb* function independently to promote L1 starvation resistance. *lin-35(n745)* and *daf-16(mu86)* null mutants were both significantly starvation sensitive compared to wild type (Fig 3A). However, the double mutant was significantly more sensitive than either single mutant, as previously reported (Cui *et al*., 2013). A two-way ANOVA suggests no interaction (additivity) between *lin-35* and *daf-16*, consistent with independent function downstream of *daf-18/PTEN*.

**Fig 3.**
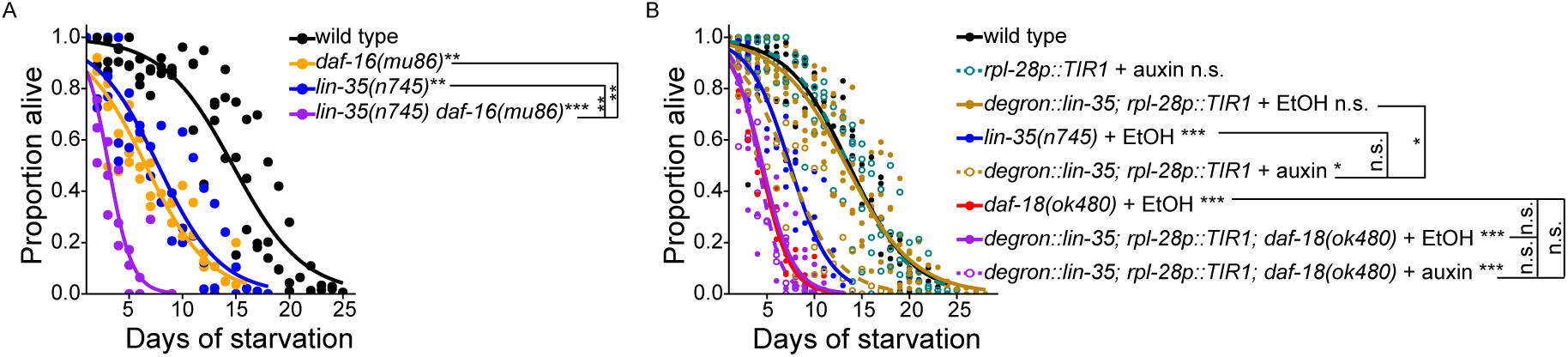
*lin-35/Rb* functions downstream of *daf-18/PTEN* to promote starvation resistance. (A and B) Proportion of survivors is plotted throughout L1 starvation. Survival was scored daily. Unless otherwise noted with brackets, all comparisons were against wild type. *P < 0.05; **P < 0.01; ***P < 0.001; n.s. not significant. For details on the statistical method, see *Statistics for starvation survival* in Materials and Methods. (A) Three biological replicates were performed. Number of scored animals was 94 +/- 28 (mean +/- standard deviation). Two-tailed, unpaired, variance-pooled t-tests were performed on half-lives to compare genotypes. Half-lives of all four genotypes were subjected to Levene’s test to assess variance homogeneity across groups, which suggested homogeneous variance. Half-lives were used in a two-way ANOVA (formula: half-life ∼ loss of *lin-35* * loss of *daf-16*) to assess additivity of the effects of *lin-35* and *daf-16* on survival, which yielded an interaction p-value of 0.1, consistent with additivity (lack of interaction or independence) of *lin-35* and *daf-16*. (B) Seven biological replicates were performed. Number of scored animals was 96 +/- 25 (mean +/- standard deviation). Two-tailed, unpaired, variance-unpooled t-tests were performed on half-lives to compare conditions. Half-lives of *degron::lin-35; rpl-28::TIR1* +/- auxin and *degron::lin-35; rpl-28::TIR1; daf-18(ok480)* +/- auxin were subjected to Levene’s test to assess variance homogeneity across groups, which suggested homogeneous variance. Half-lives were used in a two-way ANOVA (formula: half-life ∼ loss of *daf-18* * auxin addition to degrade LIN-35) to assess additivity of the effects of *lin-35* and *daf-18* on survival, which yielded an interaction p-value of 0.003, suggesting non-additivity (an interaction or dependence) of *lin-35* and *daf-18*. Auxin (indole-3-acetic acid) was used at 200 μM and was prepared in ethanol (EtOH, the solvent). The *degron::lin-35* allele used is the same as the *degron::GFP::lin-35* allele used in Fig 4 (genotype abbreviated here). EtOH (solvent alone, no auxin) was used as control.

Our RNA-seq analysis revealed a positive correlation between the *daf-16*-independent effects of *daf-18/PTEN* and *lin-35/Rb* on gene expression, and we used epistasis analysis to determine whether they function in a common pathway. We found that *lin-35(n745); daf-18(ok480)* double null mutant is inviable, so we used the auxin-induced degradation (AID) system (Zhang *et al*, 2015) and *rpl-28*-promoter-driven TIR1 to degrade degron-tagged LIN-35 protein ubiquitously (Willis *et al*, 2021). Adding auxin to *degron::lin-35; rpl-28p::TIR1* conferred starvation sensitivity indistinguishable from that of *lin-35(n745)*, suggesting that AID resulted in potent degradation of LIN-35 protein and a null phenotype (Fig 3B). However, degrading degron::LIN-35 in a *daf-18* null mutant background did not have any effect. Furthermore, a two-way ANOVA analyzing the effects of degrading degron::LIN-35 and mutating *daf-18* suggests a significant interaction (non-additivity), as if *daf-18/PTEN* depends on *lin-35/Rb*. Taken together, Fig 3 A and Fig 3B support the conclusion that *lin-35/Rb* functions downstream of *daf-18/PTEN* to promote starvation resistance and that this effect is *daf-16*-independent.

### *daf-18/PTEN* protects LIN-35/Rb from cleavage

RNA-seq suggested that *daf-18* does not affect *lin-35/Rb* transcript abundance (S1 Data), so we asked if *daf-18/PTEN* could regulate LIN-35/Rb at the protein level. We collected whole-worm lysates of *degron::GFP::lin-35; rpl-28p::TIR1* (this strain was denoted as *degron::lin-35; rpl-28p::TIR1* in Fig 3B for simplicity) in *daf-18(ok480)* and wild-type *daf-18* background, and at the same timepoint as sample collection for RNA-seq (S1 Fig A). We performed western blots with a GFP antibody, and we observed the expected size of degron::GFP::LIN-35 at ∼143 kDa (Fig 4A). For the positive control, we added auxin to the starvation culture of *degron::GFP::lin-35; rpl-28p::TIR1* (the same way as we added auxin in Fig 3B), and we observed degradation of degron::GFP::LIN-35, as expected. The band for degron::GFP::LIN-35 also appeared dimmer in the *daf-18* mutant background. We blotted against alpha-tubulin as a sample-loading control, and we quantified alpha-tubulin-normalized degron::GFP::LIN-35 band intensity (Fig 4B). The degron::GFP::LIN-35 band intensity significantly decreased compared to no auxin addition (Fig 4B). degron::GFP::LIN-35 abundance also decreased in *daf-18(ok480)* compared to the wild type, but the p-value was 0.07 (Fig 4B; see Fig 5B). Nonetheless, these results suggest that DAF-18/PTEN promotes LIN-35/Rb stability during L1 arrest.

**Fig 4.**
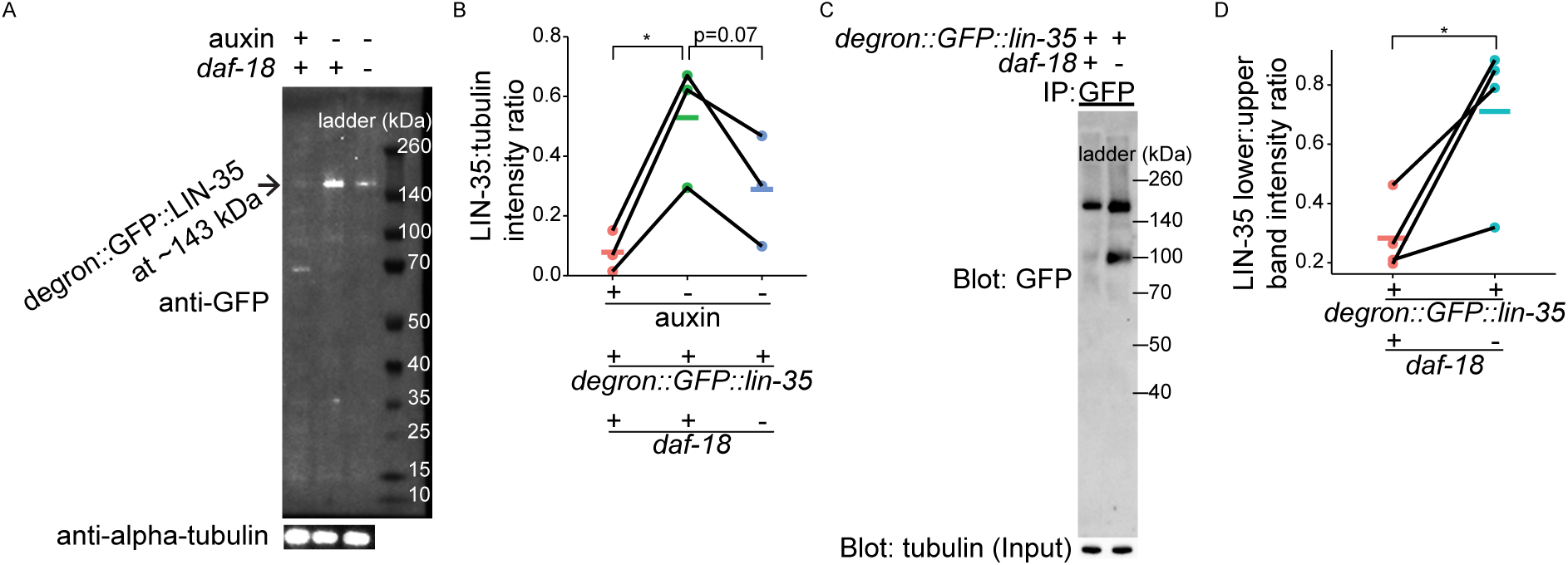
*daf-18/PTEN* protects LIN-35/Rb from cleavage. (A) Representative western blot showing the expression of LIN-35 in +/- auxin-mediated degradation and +/- *daf-18* conditions in starved L1 lysates (collected 16 hr after hypochlorite treatment) is shown. Alpha-tubulin was used as a loading control. Three biological replicates were performed (see B). (B) Quantification of LIN-35 expression level (normalized to alpha-tubulin) is plotted. Normalized LIN-35 expression levels were subjected to Bartlett’s test to assess variance homogeneity across conditions. They were then used in two-tailed, paired, variance-pooled t-tests to compare conditions, as Bartlett’s test result suggested homogeneous variance. *P < 0.05. (C) Representative western blot showing the expression of full-length and a shorter version of LIN-35 in +/- *daf-18* conditions following anti-GFP immunoprecipitation (IP) from starved L1 lysates (collected 16 hr after hypochlorite treatment). IP inputs were normalized to have the same amount of total protein, as reflected by the alpha-tubulin blot. Three biological replicates were performed (see D). (D) Quantification of the protein abundance ratio of shorter version LIN-35 and full-length LIN-35 in IP products. The ratios were subjected to Bartlett’s test to assess variance homogeneity across conditions. They were then used in two-tailed, paired, variance-pooled t-tests to compare +/- *daf-18*, as Bartlett’s test result suggested homogeneous variance. *P < 0.05. (A-D) *daf-18 (-)* refers to *daf-18(ok480)*. The *degron::GFP::lin-35* allele used here is the same as the *degron::lin-35* allele used in Fig 3.

**Fig 5.**
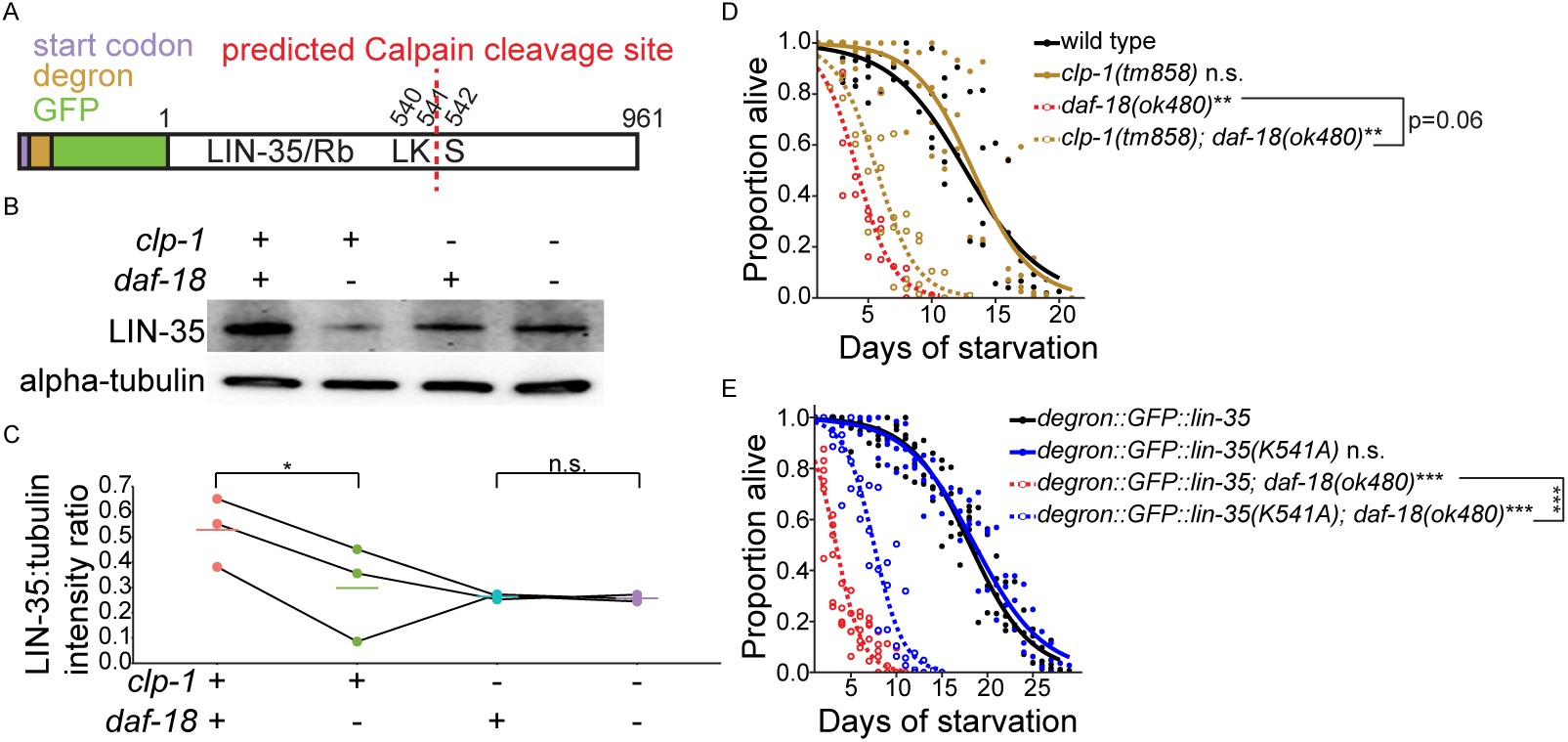
DAF-18/PTEN inhibits CLP-1/CAPN-mediated cleavage of LIN-35/Rb during starvation to promote survival. (A) A schematic of degron::GFP::LIN-35 (strain from (Willis *et al*., 2021)) is presented. A Calpain-cleavage site was predicted based on amino acid similarity to the μ-Calpain-cleavage site in human pRB (Cuerrier *et al*., 2005; Darnell *et al*., 2007; Tomita *et al*., 2020; Tompa *et al*., 2004). Indicated amino acid positions are relative to native LIN-35/Rb. (B) A representative western blot showing the expression of LIN-35 in +/- *clp-1* and +/- *daf-18* conditions in starved L1 lysates (collected 16 hr after hypochlorite treatment) is shown. Alpha-tubulin was used as a loading control. Three biological replicates were performed (see C). (C) Quantification of LIN-35 expression level (normalized to alpha-tubulin) is plotted. Normalized LIN-35 expression levels were subjected to Bartlett’s test to assess variance homogeneity across conditions. They were then used in two-tailed, paired, variance-unpooled t-tests to compare conditions, as Bartlett’s test result suggested heterogeneous variance. (B and C) *daf-18 (-)* refers to *daf-18(ok480)* and *clp-1* (-) refers to *clp-1(tm858)*. (D and E) Proportion of survivors throughout L1 starvation is plotted. Two-tailed, unpaired, variance-pooled t-tests were performed on half-lives to compare genotypes. Unless otherwise noted with brackets, all comparisons were against wild type. For details on the statistical method, see *Statistics for starvation survival* in Materials and Methods. (D) Four biological replicates were performed. Number of scored animals was 101 +/- 30 (mean +/- standard deviation). (E) Six biological replicates were performed. Number of scored animals was 121 +/- 26 (mean +/- standard deviation). (C-E) *P < 0.05; **P < 0.01; ***P < 0.001; n.s. not significant. See also S2 Fig.

We used a GFP antibody to immunoprecipitate (IP) degron::GFP::LIN-35 to investigate what happens to it in the absence of DAF-18/PTEN. We blotted with a GFP antibody, and the expected full-length degron::GFP::LIN-35 band at ∼143 kDa was present (Fig 4C). However, to our surprise, a shorter band at ∼100 kDa was also evident in the *daf-18(ok480)* background. Quantification showed that enrichment of this smaller fragment in the mutant is statistically significant (Fig 4D). Together these results suggest that DAF-18/PTEN protects LIN-35/Rb from being cleaved during L1 arrest, which produces a shorter fragment and reduces abundance of full-length LIN-35/Rb.

### DAF-18/PTEN inhibits CLP-1/CAPN-mediated cleavage of LIN-35/Rb during starvation to support survival

μ-Calpain CAPN1 cleaves pRB after lysine 810 in human cervical cancer cell lines (Darnell *et al*, 2007; Tomita *et al*, 2020), suggesting that a Calpain homolog could cleave LIN-35/Rb in the absence of DAF-18/PTEN. The closest *C. elegans* Calpain homolog to μ-Calpain is CLP-1/CAPN (S2 Fig A). We predicted a CLP-1-cleavage site after K541 in LIN-35 based on amino acid similarity to the μ-Calpain-cleavage site in human pRB (Fig 5A) (Cuerrier *et al*, 2005; Darnell *et al*., 2007; Tomita *et al*., 2020; Tompa *et al*, 2004). We subjected *degron::GFP::lin-35; rpl-28p::TIR1* with and without *daf-18(ok480)* IP lysates in Fig 4 C and D to liquid chromatography-tandem mass spectrometry (LC-MS/MS) and examined degron::GFP::LIN-35 peptide intensities in *daf-18(ok480)* vs. wild type. The results suggest an enrichment of N-terminal peptides (before K541) in *daf-18(ok480)* compared to wild type (S2 Fig B and C), which is consistent with Fig 4 C and D, since GFP was added to the N-terminus of LIN-35.

We hypothesized that CLP-1/CAPN negatively regulates LIN-35/Rb stability, so we asked if mutating *clp-1/CAPN* would rescue full-length degron::GFP::LIN-35 abundance in *daf-18(ok480)*. We collected whole-worm lysates of *degron::GFP::lin-35; rpl-28p::TIR1* with and without *daf-18(ok480)* and with and without *clp-1(tm858)* loss-of-function mutation, and we blotted against GFP. Full-length degron::GFP::LIN-35 abundance decreased in *daf-18(ok480)*, as expected (Fig 4 A and B), and this time it was statistically significant (Fig 5 B and C). Critically, this decrease in LIN-35 abundance was abolished in the *clp-1(tm858)* background (Fig 5 B and C), supporting the conclusion that CLP-1/CAPN targets LIN-35/Rb for cleavage.

Our biochemical analysis suggests that *daf-18/PTEN* mutants are starvation-sensitive in part due to decreased abundance of full-length LIN-35/Rb. We tested this hypothesis with a *lin-35* overexpression transgene (Fay *et al*, 2002), which modestly but significantly increased *daf-18(ok480)* starvation sensitivity but had no effect in a wild-type background (S2 Fig D). We sought to further test this hypothesis by protecting LIN-35 from cleavage. Specifically, we hypothesized that blocking cleavage would not have an effect in a wild-type background but would rescue starvation sensitivity of *daf-18(ok480)*. Mutating *clp-1* nearly rescued *daf-18(ok480)* starvation sensitivity (p = 0.06), but it did not make a difference in wild type (Fig 5D). We also created a point mutant that changes the predicted cleavage site lysine to alanine (K541A, in endogenous LIN-35 coordinates). A similar mutation in human *RB1* renders pRB resistant to Calpain-cleavage (Tomita *et al*., 2020). Notably, the cleavage-resistant mutant clearly and significantly rescued *daf-18(ok480)* starvation sensitivity, and it did not make a difference in wild type (Fig 5E). These results demonstrate physiological significance of CLP-1/CAPN-mediated cleavage of LIN-35/Rb, supporting the conclusion that DAF-18/PTEN prevents CLP-1/CAPN from cleaving LIN-35/Rb during L1 arrest to promote survival.

### The DREAM complex represses transcription of germline genes downstream of *daf-18/PTEN* to support starvation survival

RB family proteins can repress transcription by forming a complex with E2F/DP transcription factors, and in some cases five MuvB proteins (LIN-52, LIN-9, LIN-54, LIN-37, and LIN-53) are further recruited to form the DREAM complex (Fischer & Muller, 2017; Fischer *et al*., 2022) (Fig 6A). LIN-35/Rb shares transcriptional targets and physically interacts with DREAM components (Gal *et al*., 2021; Goetsch *et al*, 2017; Goetsch & Strome, 2022; Harrison *et al*, 2006; Latorre *et al*, 2015). Additionally, two THAP (Thanatos Associated Proteins) domain proteins are thought to mediate distinct functions of *C. elegans* DREAM, with LIN-36 mediating repression of cell cycle genes and LIN-15B mediating repression of germline genes in the soma (Gal *et al*., 2021) (Fig 6A). However, it is unclear if LIN-35/Rb functions with E2F/DP, DREAM, or either THAP domain protein in promoting starvation resistance.

**Fig 6.**
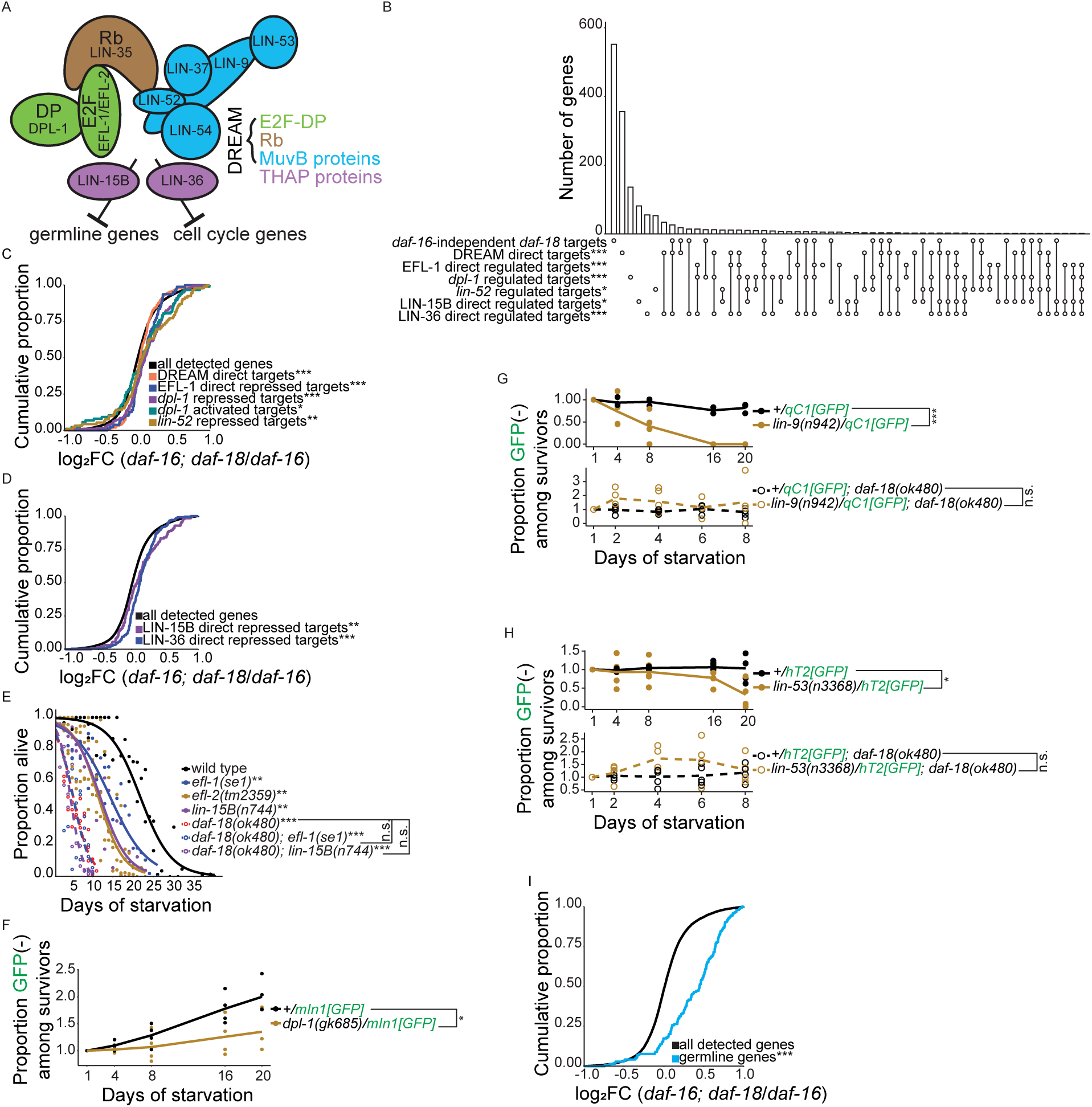
The DREAM complex represses transcription of germline genes downstream of *daf-18/PTEN* to support starvation survival. (A) A schematic of the DREAM complex and THAP-domain proteins is presented (adapted from (Gal *et al*., 2021; Goetsch & Strome, 2022)). (B) Overlaps among *daf-16*-independent *daf-18* targets (defined in Fig 2C) and targets of DREAM, E2F, MuvB, and THAP-domain proteins are plotted. Single open circles indicate the number of genes in each set that do not overlap with any other set, and vertical lines connecting open circles indicate the number of genes in the intersection of those gene sets. Targets activated and repressed by each factor are combined (based on RNA-seq analysis of mutants; “regulated”). “Direct” targets are based on ChIP-seq. Hypergeometric tests were performed to assess the overlap significance (indicated with asterisks) between *daf-16*-independent *daf-18* targets and each of the other gene sets, with the background being all detected genes in RNA-seq (see S1 Data). See C and D for published data sources. (C) Cumulative distributions of gene expression changes (log_2_FCs) in *daf-16; daf-18* vs. *daf-16* are plotted for DREAM direct target genes (n = 510; targets from (Latorre *et al*., 2015)), EFL-1 direct repressed targets (n = 79; determined by CHIP-seq and RNA-seq in (Gal *et al*., 2021)), *dpl-1* repressed and activated targets (n = 181 and n = 94, respectively; determined by RNA-seq in (Gal *et al*., 2021)), and *lin-52* repressed targets (n = 96; determined by RNA-seq in (Goetsch & Strome, 2022)). EFL-1 direct activated targets (n = 1) and *lin-52* activated targets (n = 13) were excluded because there were too few. (D) Cumulative distributions of gene expression changes in *daf-16; daf-18* vs. *daf-16* are plotted for LIN-15B and LIN-36 direct repressed targets (n = 128 and n = 245, respectively; determined by CHIP-seq and RNA-seq in (Gal *et al*., 2021)). LIN-15B and LIN-36 direct activated targets (n = 15 and n = 5, respectively) were excluded because there were too few. (C and D) Kolmogorov-Smirnov tests were used to assess the equality of two cumulative distributions. All comparisons were against “all detected genes.” (E) Proportion of survivors throughout L1 starvation is plotted. Survival was scored daily. Three biological replicates were performed. Number of animals scored was 112 +/- 24 (mean +/- standard deviation). Two-tailed, unpaired, variance-pooled t-tests were performed on half-lives to compare genotypes. Unless otherwise noted with brackets, all comparisons were against wild type. Half-lives of wild type, *efl-1(se1)*, and *lin-15B(n744)* single mutants, and corresponding *daf-18* double mutants, were subjected to Levene’s test to assess variance homogeneity across groups, which suggested homogeneous variance. Half-lives were used in two-way ANOVA (formulae: half-life ∼ loss of *daf-18* * loss of *efl-1*, half-life ∼ loss of *daf-18* * loss of *lin-15B*) to assess additivity of the effects on survival of *daf-18* and *efl-1* and also *daf-18* and *lin-15B*, which yielded an interaction p-value of 0.002 for both tests, suggesting non-additivity (an interaction or dependence) of *daf-18* with *efl-1* and *lin-15B*. (F-H) Proportion of GFP-negative worms (zygotic homozygous mutants) among survivors (normalized by Day 1) throughout L1 starvation is plotted. Parental genotypes are indicated in the legend. Survival was scored at five timepoints (days 1, 4, 8, 16, 20 for strains without *daf-18(ok480)* and days 1, 2, 4, 6, 8 for strains with *daf-18(ok480)*). *daf-18(ok480)* mutants die rapidly during starvation (Fig 1), and strains carrying this allele could not be assayed beyond 8 days. Proportion GFP(-) worms among survivors for each pair of genotypes in F-H (connected by brackets) was subjected to Levene’s test to assess variance homogeneity across groups, which suggested heterogeneous variance. Those proportions were subjected to a non-parametric two-way ANOVA (formula: proportion GFP(-) among survivors ∼ genotype * duration of starvation) using the R package ARTool to assess whether *dpl-1(gk685)* in F, *lin-9(n942)* in G, and *lin-53(n3368)* in H displayed different levels of starvation resistance than wild type. Asterisks represent p-values of the interaction between genotype and duration of starvation in non-parametric two-way ANOVA. (F) Four biological replicates were performed. Number of animals scored was 351 +/- 123 (mean +/- standard deviation). Proportion GFP(-) worms among survivors for +/*mIn1[GFP]* was subjected to Bartlett’s test to assess variance homogeneity across groups, which suggested heterogeneous variance. Those proportions were subjected to a non-parametric one-way ANOVA (formula: proportion GFP(-) among survivors ∼ duration of starvation) using the kruskal.test function in R to assess whether proportion GFP(-) worms among survivors of +/*mIn1[GFP]* changed over time, which yielded a p-value of 0.003. (G) Number of animals scored was 399 +/- 115 (mean +/- standard deviation). (H) Number of animals scored was 247 +/- 78 (mean +/- standard deviation). (G and H) Five biological replicates were performed. (I) Cumulative distributions of log_2_FCs in *daf-16; daf-18* vs. *daf-16* are plotted for a high-confidence germline gene set (Fry *et al*., 2021) (denoted “germline genes”; n = 161). (C, D, and I) Kolmogorov-Smirnov tests were used to assess the equality between two cumulative distributions. All comparisons were against “all detected genes.” (B-I) *P < 0.05; **P < 0.01; ***P < 0.001; n.s. not significant. See also S3 Fig.

Given that *daf-18/PTEN* depends on *lin-35/Rb* to promote starvation resistance, we wanted to know whether the DREAM complex or just E2F/DP is a transcriptional effector of *daf-18/PTEN.* Together with RB family proteins, E2F/DP and DREAM repress transcription (Henley & Dick, 2012; Sadasivam & DeCaprio, 2013; Uchida, 2016), and DREAM is a transcriptional repressor in *C. elegans* (Gal *et al*., 2021; Goetsch *et al*., 2017; Latorre *et al*., 2015; Petrella *et al*., 2011; Rechtsteiner *et al*., 2019), so we hypothesized that loss of *daf-18/PTEN* relieves E2F/DP or DREAM repression of *daf-18* targets independent of *daf-16/FoxO*. We revisited our RNA-seq data and found that *daf-16*-independent *daf-18* targets (defined in Fig 2C) are enriched for “EFL-1 direct regulated targets” (Gal *et al*., 2021) (Fig 6B) and that *daf-16; daf-18*/*daf-16* log_2_FCs are greater for “EFL-1 direct repressed targets” than for all detected genes (Fig 6C). Furthermore, “*dpl-1* regulated targets” displayed similar enrichment (Fig 6B), and expression of “*dpl-1* repressed targets” was increased in the double mutant (Fig 6C). Together these results suggest that the *daf-16*-independent effects of *daf-18* depend on E2F/DP. “DREAM direct targets” (bound by eight *C. elegans* DREAM components) (Latorre *et al*., 2015) were enriched among *daf-16-*independent targets of *daf-18* (Fig 6B), and their expression was also significantly greater in the *daf-16; daf-18* double mutant than the *daf-16* single mutant (Fig 6C), suggesting that the *daf-16*-independent effects of *daf-18* also depend on DREAM. The *lin-52(3A)* mutation was engineered to sever physical association between LIN-35/Rb and MuvB components of DREAM, and *lin-52(3A)-*upregulated genes (“*lin-52* repressed targets” determined from RNA-seq) represent targets of repression by an intact DREAM complex (Goetsch & Strome, 2022). *lin-52* repressed targets are also enriched among *daf-16-* independent *daf-18* targets (Fig 6B), and their expression was significantly greater in the *daf-16; daf-18* double mutant than the *daf-16* single mutant (Fig 6C), further suggesting a role of DREAM in transcriptional repression downstream of DAF-18/PTEN. However, “*dpl-1* activated targets” also had higher expression in the *daf-16; daf-18* double mutant than the *daf-16* single mutant, albeit with a smaller effect size (Fig 6C). Increased expression of *dpl-1* activated targets is seemingly inconsistent with a unitary role of DREAM in repression, but the overlap could be because they are not all direct targets (they were determined by RNA-seq without CHIP-seq) (Gal *et al*., 2021), and so they may include secondary effects of alleviating transcriptional repression on direct targets. Taken together, these results support the hypothesis that DREAM functions as a transcriptional repressor downstream of DAF-18/PTEN.

We extended our analysis to the two THAP domain proteins LIN-15B and LIN-36 (Gal *et al*., 2021). “LIN-15B direct regulated targets” and “LIN-36 direct regulated targets” are both enriched among *daf-16*-independent targets of *daf-18* (Fig 6B). In addition, both had larger *daf-16; daf-18*/*daf-16* log_2_FCs than all detected genes (Fig 6D). Notably, a negative control gene set, “*daf-18*-independent targets of *daf-16*” (defined in S1 Fig C and D), is not enriched for any of the six other gene sets included in Fig 6A (S3 Fig A). These results suggest that in addition to intact DREAM, LIN-15B and LIN-36 also contribute to transcriptional repression downstream of DAF-18/PTEN.

The preceding analysis revealed positive correlations between *daf-16*-independent effects of *daf-18* and a variety of DREAM components and mediators. We used epistasis analysis to determine whether E2F/DP, DREAM, and the THAP domain proteins function downstream of *daf-18/PTEN.* Our model predicts that mutating DREAM components on their own causes a starvation-sensitive phenotype but that they do not affect the *daf-18* null mutant, like *lin-35/Rb* mutants. Loss-of-function mutants of *C. elegans* E2F genes, *efl-1(se1)* and *efl-2(tm2359)* were starvation sensitive (Fig 6E). Furthermore, *daf-18(ok480); efl-1(se1)* was no more sensitive than *daf-18(ok480)*, and the interaction between *daf-18* and *efl-1* was significant in a two-way ANOVA (Fig 6E), suggesting that *daf-18* depends on *efl-1/E2F* to promote starvation resistance. We analyzed the null mutant *dpl-1(gk685)*, which is inviable and must be maintained with a balancer chromosome. However, the GFP-marked balancer, *mIn1[GFP]*, caused starvation sensitivity by itself, as the proportion of wild-type worms (GFP(-) progeny of +/*mIn1[GFP]*) among survivors went up significantly over time (Fig 6F). Nevertheless, two-way ANOVA (interaction between genotype and duration of starvation) suggested that *dpl-1(gk685)* is starvation sensitive compared to wild type (Fig 6F). These results reinforce the conclusion that *daf-18/PTEN* depends on E2F/DP to promote starvation resistance.

We analyzed two MuvB genes, *lin-9* and *lin-53*, to determine if *daf-18/PTEN* depends on MuvB/DREAM to promote starvation resistance. Like *dpl-1(gk685)*, the null mutants *lin-9(n942)* and *lin-53(n3368)* are inviable and were maintained with a GFP-marked balancer, except the balancers used (*qC1[GFP]* and *hT2[GFP]*) did not cause starvation sensitivity on their own (Fig 6 G and H). However, two-way ANOVA (interaction between genotype and duration of starvation) suggested that the proportion of homozygous *lin-9(n942)* and *lin-53(n3368)* worms among survivors went down over time (Fig 6 G and H), suggesting that these MuvB mutants are starvation sensitive. Critically, *lin-9(n942)* and *lin-53(n3368)* were no more sensitive in a *daf-18(ok480)* null mutant background, as suggested by two-way ANOVA (interaction between genotype and duration of starvation) (Fig 6 G and H). These results show that the positive correlations between *daf-18* and DREAM targets revealed in Fig 6 B and C are functional, supporting the conclusion that DREAM functions downstream of DAF-18/PTEN to promote starvation resistance.

We analyzed *lin-36* and *lin-15B* to determine if *daf-18/PTEN* depends on either THAP domain protein to promote starvation resistance. Interestingly, the *lin-15B(n744)* null mutant was starvation sensitive and non-additive with *daf-18(ok480)*, but *lin-36(we36)* null mutants did not affect starvation resistance in the wild-type or *daf-18(ok480)* background (S3 Fig B). These results suggest that LIN-35-DREAM promotes L1 starvation resistance downstream of DAF-18/PTEN via the THAP domain protein LIN-15B but that LIN-36 is dispensable, despite overlapping targets of LIN-36 and *daf-18* (Fig 6 B and D).

Disruption of *daf-18/PTEN*, *lin-35/Rb*, *lin-37/MuvB*, and *lin-15B/THAP* results in up-regulation of germline genes, and in the three latter cases this has been shown to be due to ectopic expression in the soma. We hypothesized that the *daf-16*-independent effects of *daf-18* (defined in Fig 2C), which are mediated at least in part through LIN-35/Rb and DREAM, subsume germline genes. We used a set of 161 “high-confidence germline genes” derived from the intersection of targets from four published germline gene sets (Fry *et al*., 2021), and found that they were significantly enriched among *daf-16*-independent targets of *daf-18* (p = 8.4 x 10^-7^). Notably, high-confidence germline genes were depleted among *daf-18*-independent targets of *daf-16* with near significance (depletion p = 0.051), suggesting a specific association between *daf-16*-independent targets of *daf-18* and germline genes. Moreover, *daf-16; daf-18/daf-16* log_2_FCs were significantly greater for high-confidence germline genes than for all detected genes (Fig 6I). These results suggest that DAF-18/PTEN acts through LIN-35/Rb and DREAM to repress germline gene expression during L1 arrest.

## DISCUSSION

Starvation resistance is intimately related to human health and disease, but the molecular basis for it is not well understood. Here we show that the tumor suppressor *daf-18/PTEN* promotes starvation resistance in *C. elegans* independent of its well-established regulation of PI3K and IIS, suggesting a critical function of DAF-18/PTEN protein-phosphatase activity (summarized in Fig 7). We discovered that DAF-18/PTEN protects another important tumor suppressor, LIN-35/Rb, from being cleaved by CLP-1/CAPN during starvation, permitting LIN-35/Rb to promote starvation resistance. LIN-35/Rb is the sole RB homolog/pocket protein in *C. elegans*, and we show that the DREAM complex and the THAP domain protein LIN-15B are required for *daf-18/PTEN* to promote starvation resistance. Our results suggest that DREAM is a transcriptional effector of DAF-18/PTEN, and that DAF-18/PTEN promotes starvation resistance through its protein-phosphatase activity by repressing expression of germline genes via DREAM.

**Fig 7.**
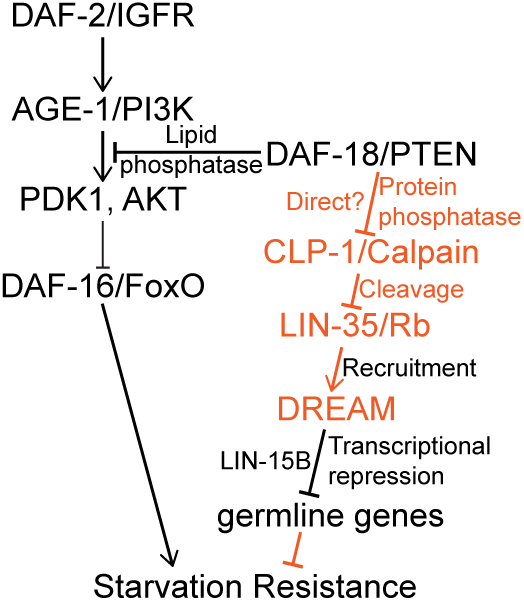
DAF-18/PTEN protein phosphatase acts through LIN-35/Rb and DREAM by inhibiting CLP-1/Calpain to promote starvation resistance. Proteins, regulatory relationships, and mechanisms known to regulate starvation resistance are in black, and those suggested by this study’s results are in orange. See Discussion for details.

### PI3K and IIS-independent effects suggest DAF-18/PTEN protein-phosphatase activity promotes starvation resistance

PTEN is a potent tumor suppressor, and *C. elegans daf-18/PTEN* promotes developmental arrest and survival during L1 starvation (Fukuyama *et al*., 2006; Fukuyama *et al*., 2012). PTEN is both a lipid (Myers *et al*., 1998) and a protein phosphatase (Myers *et al*, 1997), but its protein-phosphatase activity has not been connected to starvation resistance or maintenance of cellular quiescence. DAF-18 is the sole PTEN homolog in *C. elegans*, and DAF-18 harbors a dual-phosphatase domain like PTEN (Chen *et al*., 2022), though protein-phosphatase activity has not been demonstrated. We show that loss of *daf-18/PTEN* reduces starvation resistance in an *age-1/PI3K* null background, that gain-of-function alleles effectively increasing PI3K signaling do not phenocopy *daf-18/PTEN* null alleles, and that loss of DAF-16/FoxO, the principal effector of PI3K/IIS in this context (Baugh & Hu, 2020), does not reduce starvation resistance in a *daf-18/PTEN* null background (Fig 1). These results provide evidence that DAF-18/PTEN protein-phosphatase activity promotes starvation resistance. Furthermore, we identify LIN-35/Rb, another important tumor suppressor known to promote starvation resistance in *C. elegans* (Cui *et al*., 2013), as a mediator of PI3K/IIS-independent effects of DAF-18/PTEN (Fig 2 and 3). However, we recognize that DAF-18/PTEN may have non-phosphatase regulatory activity. For example, PTEN interacts with histone H1 via its C-terminal tail to regulate chromatin condensation independent of its phosphatase activity (Chen *et al*, 2014), and PTEN interacts with but does not change the phosphorylation state of HSD17B8 (Zhao *et al*, 2024). Nonetheless, we believe the simplest interpretation of our results together with prior knowledge is that DAF-18/PTEN lipid and protein-phosphatase activities promote starvation resistance during L1 arrest.

It is desirable to use genetic analysis to dissect the function of *daf-18/PTEN* lipid and protein-phosphatase activities, and a pair of missense mutants have been presumed to selectively disrupt these two activities (Brisbin *et al*., 2009; Nakdimon *et al*., 2012; Solari *et al*., 2005; Zheng *et al*., 2018). However, biochemical analysis of homologous mammalian PTEN mutations reveals non-selective disruption of these two activities (Furnari *et al*, 1998; Lee *et al*, 1999; Myers *et al*., 1998; Ramaswamy *et al*, 1999; Rodriguez-Escudero *et al*, 2011; Xiao *et al*, 2007). Furthermore, careful genetic analysis of these alleles engineered in *C. elegans* with genome editing revealed that they have overlapping effects on *daf-18/PTEN* function, affecting starvation resistance in L1 larvae and dauer formation (Chen *et al*., 2022; Wittes & Greenwald, 2022). Therefore, it is currently impossible to use genetic analysis to cleanly dissect the two phosphatase activities of PTEN or its homologs in regulating starvation resistance or other phenotypes.

### DAF-18/PTEN protects LIN-35/Rb from CLP-1/CAPN-mediated cleavage during starvation

Implication of DAF-18/PTEN putative protein-phosphatase activity raises the question of what protein(s) DAF-18/PTEN targets to promote starvation resistance. We used transcriptome analysis to characterize the PI3K-independent effects of *daf-18/PTEN* on gene expression to identify a candidate target. By using the expression profile from bulk RNA-seq as a phenotype for epistasis analysis between *daf-18/PTEN* and *daf-16/FoxO*, we were able to determine through multiple analyses that disruption of *daf-18/PTEN* affects gene expression independently of *daf-16/FoxO* and by extension PI3K and IIS (Fig 2). Moreover, by directly comparing a *daf-16; daf-18* double null mutant and a *daf-18* null mutant, we were able to isolate the putative effects of DAF-18/PTEN protein-phosphatase activity on gene expression during starvation. We used the resulting gene set to query a database of published results, discovering a significant positive correlation between the effects of DAF-18/PTEN protein-phosphatase activity and LIN-35/Rb on gene expression. We used epistasis analysis to confirm that *daf-16/FoxO* and *lin-35/Rb* function independently (Cui *et al*., 2013) and, critically, to demonstrate that *daf-18/PTEN* depends on *lin-35/Rb* to promote starvation resistance (Fig 3). These results suggest that *daf-18/PTEN* and *lin-35/Rb* are in a linear pathway. Furthermore, they suggested the possibility that LIN-35/Rb could be directly dephosphorylated by DAF-18/PTEN.

Multiple lines of evidence suggest that LIN-35/Rb is not a direct target of DAF-18/PTEN. We performed LC-MS/MS on starved L1 lysates after immunoprecipitating LIN-35/Rb to detect post-translational modifications in wild type and a *daf-18/PTEN* null mutant, and we did not detect an effect on LIN-35/Rb phosphorylation. We were also unable to detect LIN-35/Rb above background by LC-MS/MS or western blot after immunoprecipitating DAF-18/PTEN from a *daf-18* “trapping mutant” background (Chen *et al*., 2022; Flint *et al*, 1997). These negative results prompted us to consider alternative hypotheses for how DAF-18/PTEN promotes LIN-35/Rb activity during L1 arrest.

We found that LIN-35/Rb is destabilized in the absence of DAF-18/PTEN in starved L1 larvae (Fig 4). Furthermore, a reduced molecular weight fragment of LIN-35/Rb was evident in the absence of DAF-18/PTEN, suggesting that DAF-18/PTEN protects LIN-35/Rb from cleavage. μ-Calpain cleaves pRB in cervical cancer cells (Darnell *et al*., 2007; Tomita *et al*., 2020), so we investigated CLP-1/CAPN, the most similar *C. elegans* homolog of μ-Calpain (S2 Fig A). Multiple lines of evidence support the conclusion that DAF-18/PTEN antagonizes CLP-1/CAPN, which otherwise cleaves LIN-35/Rb to limit starvation resistance: LIN-35/Rb contains a predicted μ-Calpain cleavage site at an appropriate location to account for the size of the LIN-35/Rb cleavage product, mutation of *clp-1/CAPN* rescues cleavage of LIN-35/Rb and also starvation resistance in the absence of DAF-18/PTEN, and, critically, a point mutation disrupting the predicted μ-Calpain cleavage site in LIN-35/Rb rescues starvation resistance in a *daf-18/PTEN* null background (*i.e.*, it is a gain-of-function allele) (Fig 5). These results are consistent with DAF-18/PTEN protein-phosphatase activity promoting starvation resistance via positive regulation of LIN-35/Rb, but they suggest that this regulation is indirect. However, it is unclear if in fact DAF-18/PTEN inhibits CLP-1/CAPN directly via dephosphorylation (Fig 7).

### DREAM functions as a transcriptional effector of DAF-18/PTEN to promote starvation resistance by repressing germline gene expression

*daf-18/PTEN* represses expression of germline genes during L1 starvation, but DAF-16/FoxO is not responsible, and the transcriptional effector of DAF-18/PTEN is unknown (Fry *et al*., 2021). It has also been unclear whether DAF-18/PTEN protein-phosphatase activity is involved in transcriptional regulation. Identification of the pocket protein LIN-35/Rb as a mediator of the effects of DAF-18/PTEN on starvation resistance suggested that either E2F/DP or the DREAM complex could function as a transcriptional effector of DAF-18/PTEN. We performed meta-analysis of various functional genomics datasets interrogating regulatory targets (determined with RNA-seq) and/or binding targets (determined with ChIP-seq) of E2F/DP and MuvB/DREAM components to determine whether the PI3K/IIS-independent effects of *daf-18/PTEN* may be mediated by E2F/DP alone or the entire DREAM complex (Fig 6A-C). Multiple components of E2F/DP and the rest of DREAM appeared to bind and regulate many of the same genes affected by the putative protein-phosphatase activity of DAF-18/PTEN, suggesting that DREAM is a transcriptional effector of DAF-18/PTEN. A similar analysis also suggested that the THAP domain proteins LIN-36 and LIN-15B are involved (Fig 6D), consistent with them functioning as mediators of DREAM repression (Gal *et al*., 2021).

We used genetic analysis to determine if in fact E2F/DP, MuvB proteins (DREAM), and the THAP proteins function downstream of *daf-18/PTEN.* Several DREAM mutants have been assayed for their effects on L1 starvation resistance in *C. elegans* (Cui *et al*., 2013). *efl-1(se1)*, *dpl-1(n2994)*, and *lin-9(n112)* were assayed, but they are likely not null (Beitel *et al*, 2000; Ceol & Horvitz, 2001). Nonetheless, *efl-1(se1)* displayed starvation sensitivity, but *dpl-1(n2994)* and *lin-9(n112)* did not (Cui *et al*., 2013). This prompted us to examine a larger panel of DREAM mutants, including putative null alleles where available. We reproduced starvation sensitivity for *efl-1(se1)*, and putative null mutants *efl-2(tm2359), dpl-1(gk685)*, *lin-9(n942)*, and *lin-53(n3368)* also conferred starvation sensitivity (Fig 6E-H). These results clearly suggest that the DREAM complex (LIN-35/Rb plus E2F/DP and MuvB proteins) promotes starvation resistance. Critically, the same mutations of *efl-1/E2F*, *dpl-1/DP*, *lin-9/MuvB*, and *lin-53/MuvB* did not affect starvation resistance in a *daf-18/PTEN* null background (Fig 6E-H), indicating that DAF-18/PTEN depends on the DREAM complex to promote starvation resistance. p107 has relatively high affinity for hsLIN-52/MuvB but pRB does not (Putta *et al*, 2022), and p107 and p130 are thought to function with DREAM while pRB is thought to function with E2F/DP but not MuvB (Henley & Dick, 2012). These observations together with our results suggest that LIN-35/Rb may share functional homology with the RB-like proteins p107 and p130 rather than pRB itself.

How does transcriptional regulation by DREAM, a repressor, promote starvation resistance? DREAM is well known for its role in regulating cell cycle genes (Fischer & Muller, 2017; Fischer *et al*., 2022), but DREAM also represses germline genes in the soma of *C. elegans* (Gal *et al*., 2021; Petrella *et al*., 2011; Rechtsteiner *et al*., 2019; Wang *et al*., 2005; Wu *et al*., 2012). Notably, LIN-35/Rb and DAF-18/PTEN also repress germline genes (Fry *et al*., 2021; Petrella *et al*., 2011; Rechtsteiner *et al*., 2019; Wang *et al*., 2005; Wu *et al*., 2012). Considering the two THAP domain proteins, LIN-15B is thought to mediate DREAM repression of germline genes in the soma, and LIN-36 is thought to mediate repression of cell cycle genes (Gal *et al*., 2021). We assayed *lin-15B* and *lin-36* null mutants (Gal *et al*., 2021; Lu, 1999), and we found that *lin-15B* promotes starvation resistance (Fig 6E) but that *lin-36* does not (S3 Fig B). Furthermore, loss of *lin-15B* does not affect starvation resistance in a *daf-18/PTEN* null mutant background (Fig 6E), indicating that DAF-18/PTEN depends on LIN-15B, but not LIN-36, in addition to DREAM to promote starvation resistance. This result and the fact that DAF-18/PTEN, LIN-35/Rb, DREAM, and LIN-15B all repress germline gene expression suggest that such repression supports starvation resistance. We therefore analyzed a set of “high-confidence germline genes” (Fry *et al*., 2021), and we found that DAF-18/PTEN represses germline gene expression through its putative protein-phosphatase activity (“*daf-16*-independent targets of *daf-18*”) but not it’s putative lipid-phosphatase activity (“*daf-18*-independent targets of *daf-16*) (Fig 6I and S3C). Taken together, these results support a model in which the protein-phosphatase activity of DAF-18/PTEN protects LIN-35/Rb from CLP-1/CAPN-mediated cleavage, and that intact LIN-35/Rb recruits DREAM to germline genes for repression during starvation (Fig 7). *daf-18/PTEN* represses germline gene expression in the germ line, but the high-confidence germline genes used for analysis are not necessarily expressed exclusively in the germline (Fry *et al*., 2021). Given ectopic expression of germline genes in the soma of *lin-35/Rb, lin-37/MuvB*, and *lin-15B/THAP* (Petrella *et al*., 2011; Rechtsteiner *et al*., 2019; Wang *et al*., 2005; Wu *et al*., 2012), we propose that DREAM represses germline gene expression in the germline and soma downstream of DAF-18/PTEN during L1 arrest.

### Concluding Remarks

This work addresses molecular mechanisms of starvation resistance in an animal model, which is a fundamental, disease-relevant trait governed by a highly conserved network of genes involved in signaling and gene regulation (Baugh, 2013; Baugh & Hu, 2020). This study is significant as it reports a novel regulatory relationship between two important tumor suppressors, DAF-18/PTEN and LIN-35/Rb. Furthermore, it illustrates the critical function of these tumor suppressors along with DAF-16/FoxO beyond repressing proliferation in promoting survival of quiescence, both developmental and cellular. These insights improve understanding of the regulatory network governing starvation resistance and inform development of interventions to mitigate cancer, diabetes, and aging.

## METHODS

### Strains used in this study

Wild type N2 is from the Sternberg Lab collection.

AWR58 *lin-35(kea7[lin-35p::degron::GFP::lin-35]) I; keaSi10[rpl-28p::TIR1::mRuby::unc-54 3’UTR +Cbr-unc-119(+)] II* is from the Reinke Lab at University of Toronto.

IC166 *daf18(ok480) IV* is from the Chin-Sang Lab at Queen’s University.

JA1850 *lin-36(we36) III* is from the Ahringer Lab at University of Cambridge.

LRB447 *lin-35 (n745) I* is from the Fay Lab at University of Wyoming and was a five-time backcrossed version of MT10430 *lin-35 (n745) I*.

BQ1 *akt-1(mg306) V*, CF1038 *daf-16(mu86) I*, GR1310 *akt-1(mg144) V*, GR1318 *pdk-1(mg142) X*, JJ1549 *efl-1(se1) V*, MT15107 *lin-53(n3368) I/hT2 [bli-4(e937) let-?(q782) qIs48] (I;III)*, MT2495 *lin-15B(n744) X* and VC1523 *dpl-1(gk685)/mIn1 [mIs14 dpy-10(e128)] II* are from the Caenorhabditis Genetics Center (CGC).

FX00858 *clp-1(tm858) III* and FX02359 *efl-2(tm2359) II* are from the National BioResource Project (NBRP)::C.elegans in Japan.

### Strains generated in this study

PHX8778 *lin-35(kea7[lin-35p::degron::GFP::lin-35] syb8778[K541A]) I* was generated by SunyBiotech.

LRB378 *daf-16(mu86) I; daf-18(ok480) IV*

LRB429 *akt-1(mg144) V; pdk-1(mg142) X*

LRB430 *daf-18(ok480) IV; akt-1(mg306) V*

LRB441 *age-1(m333) II; daf-18(ok480) IV*

LRB461 *lin-35(n745) daf-16(mu86) I*

LRB486 *lin-35(kea7[lin-35p::degron::GFP::lin-35]) I; keaSi10[rpl-28p::TIR1::mRuby::unc-54 3’UTR +Cbr-unc-119(+)] II; daf-18(ok480) IV*

LRB487 *keaSi10 [rpl-28p::TIR1::mRuby::unc-54 3’UTR + Cbr-unc-119(+)] II*

LRB548 *lin-35(kea7[lin-35p::degron::GFP::lin-35]) I*

LRB550 *lin-35(kea7) I; daf-18(syb1659[daf-18::degron::3xFLAG] syb6062[D137A]) IV*

LRB565 *+/qC1 [dpy-19(e1259) glp-1(q339)] nIs189 III*

LRB589 *+/qC1 [dpy-19(e1259) glp-1(q339)] nIs189 III; daf-18(ok480) IV*

LRB595 *clp-1(tm858) III; daf-18(ok480) IV*

LRB596 *+/hT2 [bli-4(e937) let-?(q782) qIs48] (I;III)*

LRB598 *lin-35(kea7[lin-35p::degron::GFP::lin-35]) I; keaSi10[rpl-28p::TIR1::mRuby::unc-54 3’UTR +Cbr-unc-119(+)] II; clp-1(tm858) III; daf-18(ok480) IV*

LRB599 *lin-35(kea7[lin-35p::degron::GFP::lin-35]) I; keaSi10[rpl-28p::TIR1::mRuby::unc-54 3’UTR +Cbr-unc-119(+)] II; clp-1(tm858) III*

LRB610 *kuEx119[sur-5::GFP + C32F10]*

LRB611 *daf-18(ok480) IV; kuEx119[sur-5::GFP + C32F10]*

LRB634 *daf-18(ok480) IV; efl-1(se1) V*

LRB635 *daf-18(ok480) IV; lin-15B(n744) X*

LRB638 *lin-36(we36) III; daf-18(ok480) IV*

LRB652 *lin-35(kea7[lin-35p::degron::GFP::lin-35] syb8778[K541A]) I; daf-18(ok480) IV*

LRB654 *+/hT2 [bli-4(e937) let-?(q782) qIs48] (I;III); daf-18(ok480) IV*

LRB659 *lin-35(kea7[lin-35p::degron::GFP::lin-35]) I; daf-18(ok480) IV*

LRB673 *lin-9(n942)/qC1 [dpy-19(e1259) glp-1(q339)] nIs189 III*

LRB682 *lin-9(n942)/qC1 [dpy-19(e1259) glp-1(q339)] nIs189 III; daf-18(ok480) IV*

LRB683 *+/mIn1 [mIs14 dpy-10(e128)] II*

LRB685 *lin-53(n3368) I/hT2 [bli-4(e937) let-?(q782) qIs48] (I;III); daf-18(ok480) IV*

### *C. elegans* maintenance

All strains assayed in this study were maintained with *Escherichia coli* (*E. coli)* OP50 on nematode growth medium (NGM) plates and were well-fed for at least three generations before being used in experiments. Worms were cultured and starved at 20°C. Unless otherwise noted, all processes in Materials and Methods were done at 20°C.

### Auxin preparation and addition

A 400 mM indole-3-acetic acid (auxin) master stock was prepared in ethanol and stored in the dark at -20°C. A working stock of 133 mM auxin was prepared by diluting the master stock with ethanol, and it was also stored in the dark at -20°C. For experiments, the working stock was added to the culture at 200 μM, making it 0.15% ethanol. 0.15% ethanol without auxin was added to control cultures.

### Starvation survival

For each biological replicate and each strain, seven L4 worms were picked onto four 10 cm NGM plates seeded with OP50 (total: 28 L4 larvae). 96 hr later, those large plates were hypochlorite treated to collect embryos from gravid adults (Hibshman *et al*, 2021). Those embryos were then resuspended, washed, counted, and cultured in either S-basal with 0.1% ethanol (for Fig 1, Fig 3A, Fig 5D, Fig 6D, S2 Fig D, and S3 Fig B), S-basal with 0.15% ethanol or 200 μM auxin (for Fig 3B), or S-basal with 0.15% ethanol (for Fig 5E). Cultures had ∼1 embryo/μL in 5 ml and were placed in 16 mm glass tubes at 20°C in the dark on a tissue-culture roller drum at ∼30 rpm so embryos hatched and entered L1 arrest (Hibshman *et al*., 2021). The day after hypochlorite treatment, and again every day after that, a 100 μL aliquot of each starvation culture was plated on a 6 cm NGM plate around an *E. coli* OP50 lawn in the center. The number of larvae in the aliquot was recorded (total plated). Two days later, the number of live worms on the lawn was recorded (total alive). Proportion alive was determined as total alive divided by total plated. For all strains harboring *lin-35(n745)*, *efl-1(se1)*, *lin-15B(n744)*, and the *hT2[GFP]* balancer, ten L4 larvae were picked onto at least six 10 cm NGM plates (total: at least 60 L4s), and 120 hr later, those large plates were hypochlorite treated. Everything else was the same as described above.

### Statistics for starvation survival

For each genotype in each replicate, proportion alive on each day was normalized to the first day of starvation. Survival curves were fit using quasi-binomial logistic regression, with the response variable being normalized proportion alive and the explanatory variable being days of starvation. Half-lives were calculated for each survival curve, and replicate half-lives were subjected to Bartlett’s test to assess variance homogeneity across genotypes. Two-tailed unpaired t-tests were performed on half-lives to compare genotypes, and variance was pooled if Bartlett’s test suggested homogeneous variance.

### Hatching curve for RNA-seq

Gravid adult worms were hypochlorite treated to collect embryos (see above), which were then resuspended, washed, counted, and cultured in S-basal with 0.1% ethanol. Cultures had ∼1 embryo/μL in 20 ml (∼20,000 embryos per culture) and were placed in Erlenmeyer flasks at 20°C in a shaking incubator at 180 rpm, so embryos hatched and entered L1 arrest (Hibshman *et al*., 2021). Starting 12 hr after hypochlorite treatment, a 100 μL aliquot was sampled from the culture every hour, and the numbers of hatched and unhatched embryos were recorded. Proportion hatched was calculated as the number of hatched embryos divided by the total number of sampled embryos. At 12 hr after hypochlorite treatment, the average hatching efficiency (proportion hatched) for all cultures was about 50%, and at 16 h after hypochlorite treatment, all genotypes reached maximal hatching efficiency (S1 Fig A). 16 hr after hypochlorite treatment was chosen as the timepoint for RNA-seq sample collection.

### RNA-seq sample collection

Worms were hypochlorite-treated and embryos were cultured as described in *Hatching curve for RNA-seq.* 16 hr after hypochlorite treatment, cultures were transferred to 15 mL conical tubes and spun at 3,000 rpm for 1 min. Starved larvae were transferred to 1.5 ml Eppendorf tubes with Pasteur pipets in 100 μL or less, and the tubes were snap-frozen in liquid nitrogen and stored at -80°C until RNA isolation. Three biological replicates were performed.

### RNA isolation and RNA-seq library preparation

RNA was extracted using TRIzol Reagent (Invitrogen# 15596026) using the manufacturer’s protocol with some exceptions. 100 uL of acid-washed sand (Sigma-Aldrich# 27439) was added to each sample at the beginning of the extraction protocol to aid with homogenization. RNA was eluted in nuclease-free water and stored at -80C until further use. Libraries were prepared for sequencing using the NEBNext Ultra II RNA Library Prep Kit for Illumina (New England Biolabs# E7775) starting with 50 ng of total RNA per sample as input and 14 cycles of PCR. Barcoded libraries were pooled and sequenced on the NovaSeq 6000 S-Prime flow cell to obtain 50 bp paired-end reads. See the README sheet in S1 Data for the number of reads obtained per library.

### Differential expression analysis of RNA-seq data

bowtie2-2.3.3.1-linux-x86_64 (Langmead & Salzberg, 2012) was used to map paired-end reads with the command bowtie2 -p 2 -k 1 -S.1 -m 2 -S -p 2. The average mapping efficiency was 90.0%, and the standard deviation was 1.5% (S1 Data). HTSeq python-htseq 0.6.1p1-4build1 (amd64 binary) in ubuntu bionic (Anders *et al*, 2015) was used to count reads against *C. elegans* genome version WS273. Count data was restricted to include only protein-coding genes (20,127). Before differential expression analysis, the gene list was further restricted to include only genes with counts per million (CPM) > 1 in at least three libraries (15,018 genes). Counts were then normalized using the Trimmed Mean of M-values (TMM) method using edgeR 3.28.1 (Robinson *et al*, 2010). Principal components analysis (PCA) in S1 Fig B was performed on the set of 15,018 reproducibly detected genes using prcomp function in R stats package. The glmQLFit and glmQLFTest functions in edgeR found 871 differentially expressed genes out of 15,018 in at least one genotype amongst *daf-16*, *daf-18*, *daf-16; daf-18*, and wild type. For hierarchical clustering in Fig 2A, CPM values for these 871 genes were averaged across replicates within genotype, z-score normalized, and clustered using hclust function in R stats package. The exactTest function in edgeR was used to find genes differentially expressed between pairs of genotypes. CPM values for each gene, genotype, and replicate along with p-values for differential expression analysis are available in S1 Data.

### Transcriptome-wide epistasis analysis

The transcriptome-wide epistasis coefficient is described elsewhere (Angeles-Albores *et al*., 2018), but it is essentially the slope of a regression over all significant differentially expressed genes (DEGs) when the observed log_2_ fold changes (log_2_FCs) in the double mutant is plotted as a function of the expected log_2_FCs in the double mutant based on log_2_FCs in each single mutant assuming the two genes interact log-additively (*i.e.*, function independently). Raw count values were processed and genes were filtered as described in *Differential expression analysis of RNA-seq data* (restricted to protein-coding genes and CPM > 1 in at least 3 libraries), resulting in 15,018 genes. Count normalization and differential expression analysis of each mutant vs wild type was conducted using DESeq2 1.30.1 (Love *et al*, 2014). DESeq2 results were used in a transcriptome-wide epistasis analysis pipeline as described (Angeles-Albores *et al*., 2018). Specifically, DESeq2 generated gene-wise log_2_FCs for *daf-16*, *daf-18*, and *daf-16; daf-18* compared to wild type, as well as standard errors and q-values of those log_2_FCs. Q-value < 0.1 was used as the cutoff to extract DEGs in mutants compared to wild type. The transcriptome-wide epistasis analysis pipeline (Angeles-Albores *et al*., 2018) used DEGs shared by all three mutant-vs-wild type comparisons (563 genes) to fit predefined models and a parameter-free model. The predefined models are:

- Linear/activation model: *daf-18* ➔ *daf-16*➔ expression profile
- Linear/suppression model: *daf-18* —| *daf-16* ➔ expression profile
- Log-additive model: 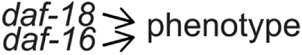

The parameter-free model does not have underlying presumptions.

Log_2_FCs for the 563 shared DEGs were bootstrapped 5,000 times (choosing 563 genes with replacement) to fit each model, and the distribution of transcriptome-wide epistasis coefficients was plotted (Fig 2B). Besides generating the distribution of transcriptome-wide epistasis coefficients, model-fitting also generated a likelihood for each model using Bayesian statistics (Angeles-Albores *et al*., 2018). Odds ratio (OR) was computed by dividing the model likelihood of the parameter-free model by each predefined model. OR > 10^3^ was used as the cutoff to reject predefined models.

### Gene set enrichment analysis

*daf-16*-independent targets of *daf-18* (as defined in Fig 2C) were analyzed with WormExp v2.0 (Yang *et al*., 2016), a web application that tests if a user’s input gene set is enriched for other experimental gene sets using Fisher’s Exact test statistics. WormExp’s default FDR < 0.1 was used as the cutoff to assess significant enrichment. The results are in S2 Data.

Hypergeometric tests were conducted for Fig 2C, 2D, 2F, 6C, S1C, S1D, and S3A. In those Venn-diagram style plots, hypergeometric tests were done by comparing two gene sets in the Venn diagram, with the background set being all genes detected by RNA-seq (n = 15,018). The other type of plot was generated using UpSetR 1.4.0 (Conway *et al*, 2017), which displays the number of genes shared by different combinations of multiple gene sets. Hypergeometric tests were done by comparing *daf-16*-independent targets of *daf-18* (Fig 6C) or *daf-18*-independent targets of *daf-16* (S3 Fig A) to the other gene sets in the plot.

Kolmogorov-Smirnov tests were used to assess the equality between cumulative distributions in Fig 2E, 2G, 6A, and 6B. All comparisons were against “all detected genes.”

### Western blot sample collection

Samples were collected the same way as described in *RNA-seq sample collection* except that starvation media were either virgin S-basal with 0.15% ethanol or 200 μM auxin, and protease inhibitors (Millipore Sigma 4693159001) were included. Frozen samples were freeze-thawed three times, using liquid nitrogen and a 45°C water bath. Then, sample buffer (Millipore Sigma S3401) was added, and samples were boiled for 10 min at 95°C, frozen for 15 min on dry ice, boiled again for 10 min at 95°C, and centrifuged at 14,000 g for 10 min to pellet any worm debris. The supernatant was collected, used to measure protein concentrations using the Pierce 660 nm Protein Assay (Thermo Fisher 22662), and used in western blots. Three biological replicates were performed.

### Western blot

∼2 μg of protein was loaded per lane on a 4-12% Bis-Tris gel (Thermo Fisher NP0321BOX). The gel was run at 200 V for 50 min and was then transferred at 30 V for 1 hr to a polyvinylidene fluoride (PVDF) membrane (Thermo Fisher LC2005). The membrane was blocked for 1 hr using 3% milk (Bio-Rad 1706404) dissolved in 1x Tris-Buffered Saline with Tween 20 (TBST), thoroughly washed three times using 1x TBST, and then incubated overnight at 4°C in 1:5,000 1x TBST-diluted primary antibody against GFP (Santa Cruz sc-9996). The membrane was then thoroughly washed three times using 1x TBST and incubated for 1 hr in 1:10,000 1x TBST-diluted horseradish peroxidase (HRP)-conjugated secondary antibody against Mouse IgG (Jackson ImmunoResearch 115-035-166). The membrane was then thoroughly washed three times using 1x TBST and then blotted using a chemiluminescent assay (Thermo Fisher 34094). Western blot images were acquired when a single-dot saturation was seen in any band on the membrane. Band intensity was quantified using ImageJ 1.54f (Schindelin *et al*, 2012) with background intensity subtracted. Western blot images and quantification are in Fig 4A, 4B, 5B, and 5C.

### Immunoprecipitation (IP) and mass spectrometry (MS) sample collection

∼6 million starved L1s were collected following the protocol in (Hibshman *et al*., 2021) with minor modifications. Specifically, gravid worms raised on plates were hypochlorite-treated to obtain embryos, and embryos were cultured in liquid at 20°C, 180 rpm, at a density of 5 worm/μL, and with 50 mg/mL *E. coli* HB101 until they were gravid adults (72 hr). Adults were washed and hypochlorite-treated to obtain millions of embryos. Those embryos were cultured in virgin S-basal with 0.1% ethanol at 5 embryos/μL in 2.8 L Erlenmeyer flasks in a shaking incubator at 180 rpm, so embryos hatched and entered L1 arrest (Hibshman *et al*., 2021). 16 hr later, those cultures were washed and collected in IP Buffer (50 mM Tris-Cl pH 7.5, 100 mM KCl, 2.5 mM MgCl_2_, 0.1% NP-40 Alternative (Millipore Sigma 492016)), which was pre-cooled to 4°C and contained phosphatase inhibitors (Millipore Sigma 4906845001) and protease inhibitors (Millipore Sigma 4693159001). At this point, samples looked slurry-like. The worm slurries were frozen into “popcorn” by dripping them into liquid nitrogen. Worm popcorn was stored at -80°C until sample processing. All sample processing was performed at 4°C. Worm popcorn was homogenized using mortar and pestles that were pre-cooled using liquid nitrogen, and the homogenate was centrifuged at 16,000 g for 10 min to remove insoluble material. The supernatant was the “input sample”. 80 μL of the input sample was used to measure protein concentrations, as described in *Western blot sample collection*. The remainder of the input sample was then diluted to 1 μg/μL with IP buffer and was used for IP.

### Immunoprecipitation (IP) and mass spectrometry (MS)

The entire IP process was done at 4°C. After diluting input samples to 1 μg/μL as described in *Immunoprecipitation (IP) and mass spectrometry (MS) sample collection*, 20 μL of anti-GFP bead slurry (ChromoTek gta) was added to each sample. The samples were incubated on a nutator for 1 hr. Then, beads were pelleted by centrifuging at 500 g for 30 sec. Beads were then washed thoroughly eight times by adding IP buffer, incubating on a nutator, pelleting beads at 500 g for 30 sec, and removing the supernatant. The total time of eight washes was ∼30 min. 70 μL of each sample was submitted to Duke Proteomics Core for MS analysis, and the remainder was used to determine sample concentrations and western blot (Fig 4 C and D), as described in *Western blot sample collection* and *Western blot*, respectively. Quantitative LC-MS/MS was performed using an MClass UPLC system (Waters Corp) coupled to a Thermo Orbitrap Fusion Lumos high-resolution accurate mass tandem Mass Spectrometer (Thermo) equipped with a FAIMSPro device via a nanoelectrospray ionization source. For additional details regarding LC-MS/MS contact the corresponding author.

## DATA AVAILABILITY

RNA-seq data: Gene Expression Omnibus GSE281157 (https://www.ncbi.nlm.nih.gov/geo/query/acc.cgi?acc=GSE281157, token cfyrgiiovdmztih)

Code to reproduce all results presented in this paper: https://github.com/jc271828/Chen2024

S1 Data: RNA-seq analysis results

S2 Data: WormExp results

S1 File: Raw images of Western Blots

## ACKNOWLEDGEMENTS

We would like to thank Aaron Reinke and David Fay for sharing strains. Some strains were provided by the CGC, which is funded by NIH Office of Research Infrastructure Programs (P40 OD010440). We would also like to thank WormBase. This work was funded by the National Institutes of Health (R01GM117408 and R01GM143159, awarded to LRB).

---

**S1 Fig.**
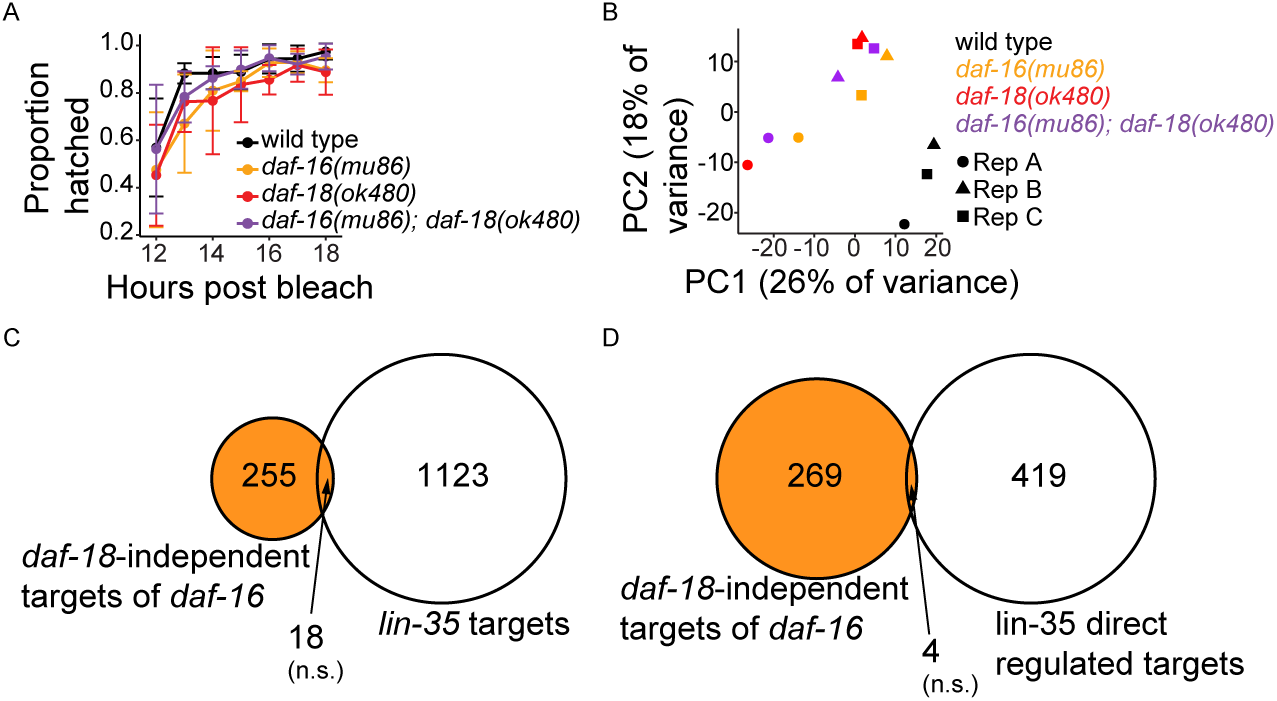
RNA-seq quality control and negative controls for gene set enrichment analysis. Related to Fig 2. (A) Proportion of hatched embryos (hatching efficiency) was measured between 12 and 18 hr after hypochlorite treatment. Four biological replicates were performed. Dots are average hatching efficiency across replicates. Error bars represent standard deviation. At ∼16 hr after bleach, all genotypes reached maximal hatching efficiency, which was chosen as the timepoint for RNA-seq sample collection. For details on how hatching efficiency was assayed, see *Hatching efficiency for determining RNA-seq sample collection timepoint* in Materials and Methods. (B) Principal-component (PC) analysis of RNA-seq data from three biological replicates (Rep A, Rep B, and Rep C). (C) Overlap between *daf-18*-independent targets of *daf-16* and *lin-35/Rb* targets in L1 arrest (from expression analysis in (Cui *et al*., 2013)). Compare to Fig 2D. (D) Overlap between *daf-18*-independent targets of *daf-16* and *lin-35* direct regulated targets in L1 arrest (determined by CHIP-seq and RNA-seq in (Gal *et al*., 2021); *i.e.*, genes bound by LIN-35 (direct) and whose expression was affected in the *lin-35* mutant (regulated)). Compare to Fig 2F. (C and D) The orange region represents *daf-18*-independent targets of *daf-16*. Hypergeometric tests were performed to assess the overlap significance between two gene sets, with the background being all detected genes in RNA-seq (see S1 Data). n.s. not significant.

**S2 Fig.**
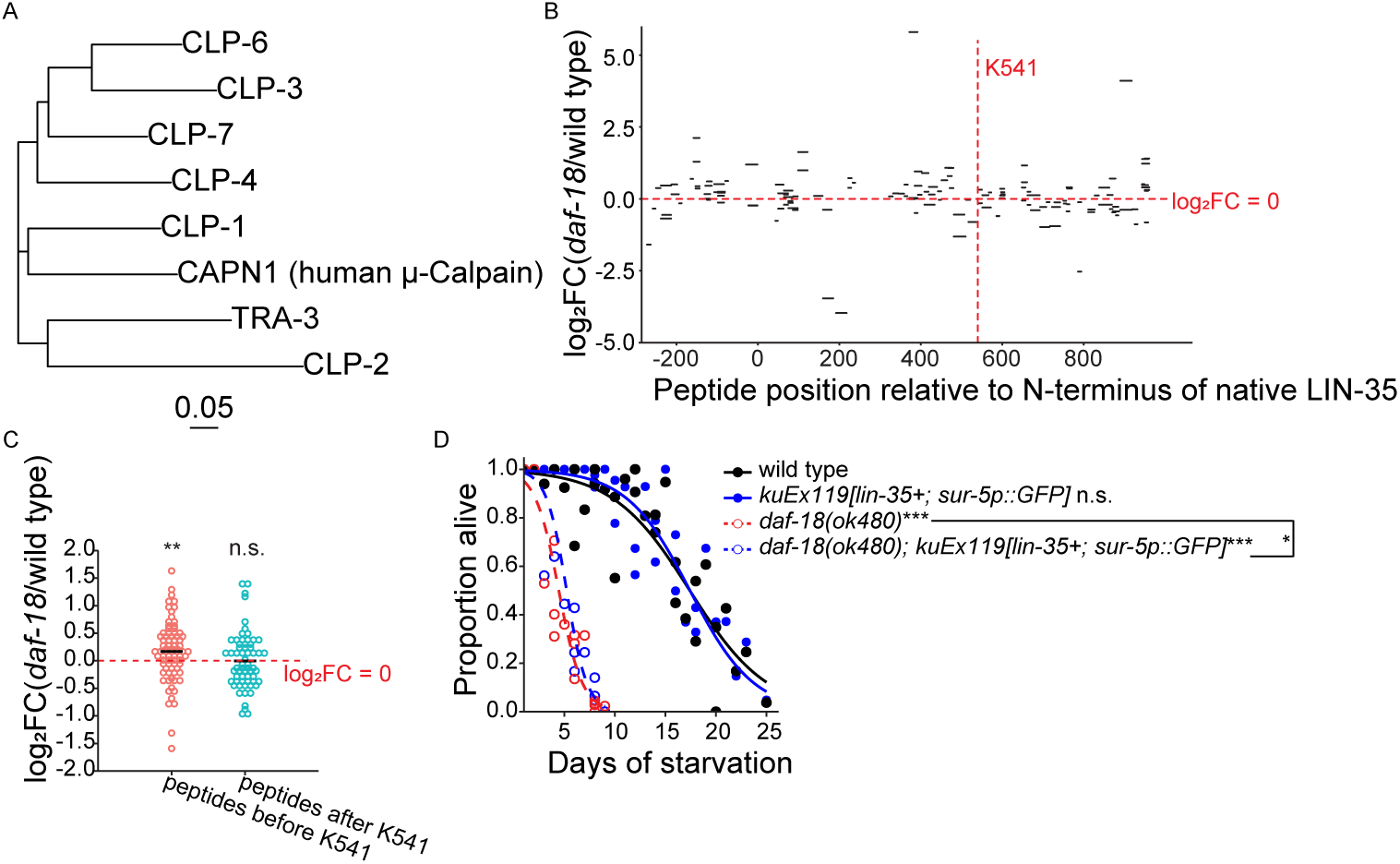
Full-length LIN-35/Rb promotes starvation resistance. Related to Fig 5. (A) Phylogenetic analysis of *C. elegans* Calpain homolog proteins and CAPN1 (human μ-Calpain). Multisequence alignment was performed using Clustal Omega (Sievers & Higgins, 2018) and plotted in R. Scale bar represents 0.05 amino acid substitutions per site. (B and C) Mass-spectrometry-based analysis of anti-GFP immunoprecipitation from starved L1 *degron::GFP::lin-35* lysates. Three biological replicates were performed. (B) log_2_ Fold Change (log_2_FC) of degron::GFP::LIN-35 peptide intensity in *daf-18(ok480)* vs. wild type plotted along native LIN-35 protein coordinates. Horizontal dashed red line indicates no enrichment (log_2_FC = 0). Vertical dashed red line indicates predicted cleavage site. (C) log_2_FC of degron::GFP::LIN-35 peptide intensity in *daf-18(ok480)* vs. wild type before and after the predicted cleavage site (K541 refers to native LIN-35/Rb). Shapiro tests were used to assess data normality. Because data was not normally distributed, two-tailed non-parametric Wilcoxon tests were used to compare log*_2_*FC to zero. (D) Proportion of survivors throughout L1 starvation. Survival was scored daily. Four biological replicates were performed. Number of scored animals was 114 +/- 30 (mean +/- standard deviation). Two-tailed, unpaired, variance-pooled t-tests were performed on half-lives to compare genotypes. Unless otherwise noted with brackets, all comparisons were against wild type. For details on the statistical method, see *Statistics for starvation survival* in Materials and Methods. (C and D) *P < 0.05; **P < 0.01; ***P < 0.001; n.s. not significant.

**S3 Fig.**
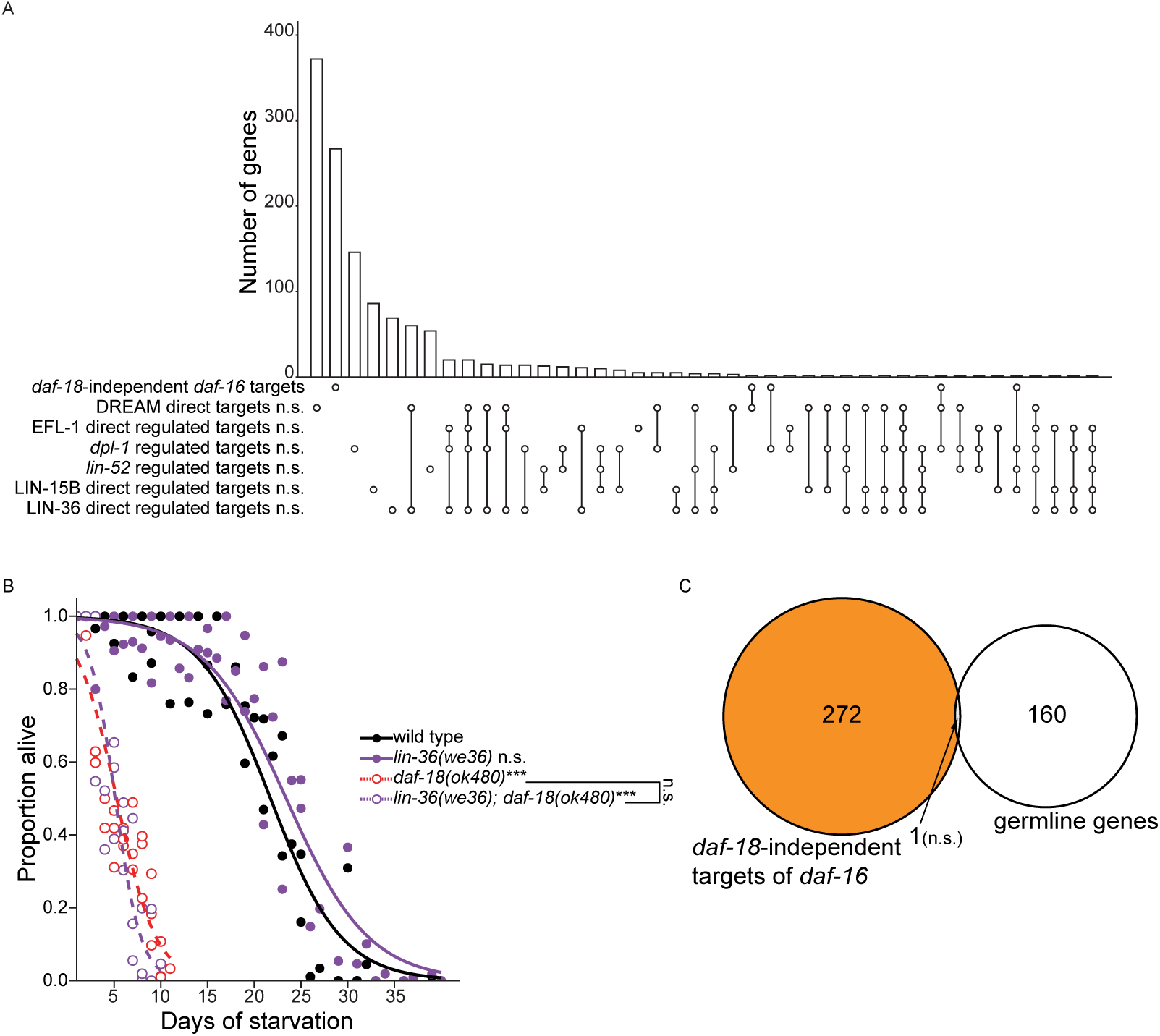
Negative controls for DREAM enrichment analyses. Related to Fig 6. (A) Overlap among gene sets in Fig 6A (included both activated and repressed targets) and *daf-18*-independent *daf-16* targets (orange regions in S1 Fig C and D) is plotted. Hypergeometric tests were performed to assess the overlap significance (indicated with asterisks) between *daf-18*-independent *daf-16* targets and each of the gene sets, with the background being all detected genes in RNA-seq (see S1 Data). Compare to Fig 6C. (B) Proportion of survivors throughout L1 starvation. Data came from the same experimental trials and were produced and analyzed the same way as in Fig 6D. Unless otherwise noted with brackets, all comparisons were against wild type. (A and B) ***P < 0.001; n.s. not significant. (C) A Venn diagram is plotted for the overlap between “high-confidence germline genes” and “*daf-18*-independent targets of *daf-16”.* The indicated enrichment p-value was calculated based on the hypergeometric distribution. The “depletion p-value” mentioned in Results equals 1 minus the enrichment p-value.

